# Distinguishing near- versus off-critical phase behaviors of intrinsically disordered proteins

**DOI:** 10.64898/2025.12.01.691696

**Authors:** Gaurav Mitra, Souradeep Ghosh, Kiersten M. Ruff, Ruoyao Zhang, Gaurav Chauhan, Rohit V. Pappu

**Affiliations:** Department of Biomedical Engineering and Center for Biomolecular Condensates, Washington University in St. Louis, St. Louis, MO, USA; Department of Chemical Engineering, Indian Institute of Technology, Indore, India

**Keywords:** critical points, universality, intrinsically disordered proteins, phase separation, percolation

## Abstract

Intrinsically disordered prion-like low complexity domains (PLCDs) drive phase transitions that underlie the biogenesis of different biomolecular condensates. The mapping of critical points is essential for generating quantitative assessments of driving forces and for distinguishing phase behaviors in the critical versus off-critical regimes. Computations play an important role in connecting the molecular-scale interactions to mesoscale phase behaviors of PLCDs and other intrinsically disordered proteins. We report results from accurate mapping of the critical regime for an archetypal PLCD. This is achieved by combining large-scale simulations with computations of Binder cumulants and the use of rigorous finite-size scaling approaches. The computed binodal, defined by knowledge of the critical point and intersection of the left arm of the binodal by the overlap and percolation lines, can be demarcated into three distinct regimes. Regime I is farthest from the critical point, with the coexisting dilute phase being akin to a gas of dispersed polymers. Regime II lies above the intersection of the overlap line and the dilute arm of the binodal. Here, coexisting dilute phases are characterized by heterogeneous cluster-size distributions with heavy tails. In Regimes I and II, dense phases are confined percolated networks. Regime III is closest to the critical point. Here, the dense phase becomes unconfined and the percolated network swells to become system-spanning. Thus, Regime III comprises two interconnected, system-spanning networks. In addition to accurately mapping the critical point, we also evaluated methods for identifying the theta temperature. We find that conventional scaling approaches lead to erroneously low estimates of the theta temperature. Instead, accurate estimation of the theta temperature requires direct calculation of the temperature dependence of the two-body interaction coefficient. We discuss the broader implications of these findings for inferring solvent quality from scaling analyses.

## Introduction

Biochemical reactions in living cells are organized in space and time by the reversible formation of membraneless bodies known as biomolecular condensates (1, 2). Phase transitions drive condensate formation (3–5) and multivalence is a defining hallmark of proteins that drive condensation (1, 5, 6). For proteins, multivalence comes in different flavors (7) and this includes: multiple protein-protein interaction domains tethered by intrinsically disordered linkers (5, 8–10), oligomeric proteins comprising folded domains with or without intrinsically disordered regions (IDRs) (11–15), or intrinsically disordered proteins (IDPs) featuring multiple cohesive motifs that provide specificity of associations (16–20).

Phase separation coupled to percolation (PSCP) is the process that explains the complete set of known phase transitions that underlie condensate formation (5, 6, 8, 9, 21–25). In this process, multivalence of specific motifs or domains enables reversible associations that lead to the formation of clusters of different sizes in solution. As concentrations increase above a threshold known as the gel point or percolation threshold (*c*_perc_), there is a finite probability of forming clusters that can grow to become system-spanning. However, at temperatures below the critical temperature *T*_c_, the growth of clusters in solution is typically suppressed because the system crosses the solubility limit (26–30). Hence, the overall free energy of the system is minimized by separation of the entire solution into coexisting dense and dilute phases via a process known as phase separation (6, 31). For systems where the associations are homotypic, the threshold concentration for phase separation is the saturation concentration or *c*_sat_. If a blend of homotypic and heterotypic associations is involved, then the solubility limit is defined by the solubility product (32–34), with different components contributing differently (34, 35). For condensation via PSCP, the macromolecular concentration within dense phases will be above *c*_perc_ for temperatures away from the critical point, and the dense phase will be a percolated network that resembles a confined physical gel (9, 23, 29, 36, 37).

It has been proposed that condensates in living cells have the potential to be in the vicinity of thermodynamic critical points (38–45). Being close to the critical point would imply that condensates behave like systems that are on the edge of stability, characterized by large fluctuations of the order parameter. For condensates, the relevant order parameters are two-fold: they are the density differences of the condensate-driving macromolecules and solution components (31) across coexisting phases, and the degree of clustering and networking of molecules (46). Importantly, the one-phase regimes for temperatures near the critical point versus far away from the critical point are expected to be very different (41, 47). Thus, the properties of the coexisting dilute phases are likely to be useful and measurable signatures of near-versus off-critical behaviors of condensates (48, 49). To set up the relevant expectations for a one-component system, we turned to simulations and mapped the complete phase behavior of A1-LCD, which is an archetypal IDP. This is the intrinsically disordered PLCD from the protein hnRNP-A1 (Fig. 1A) (50).

**Fig. 1.**
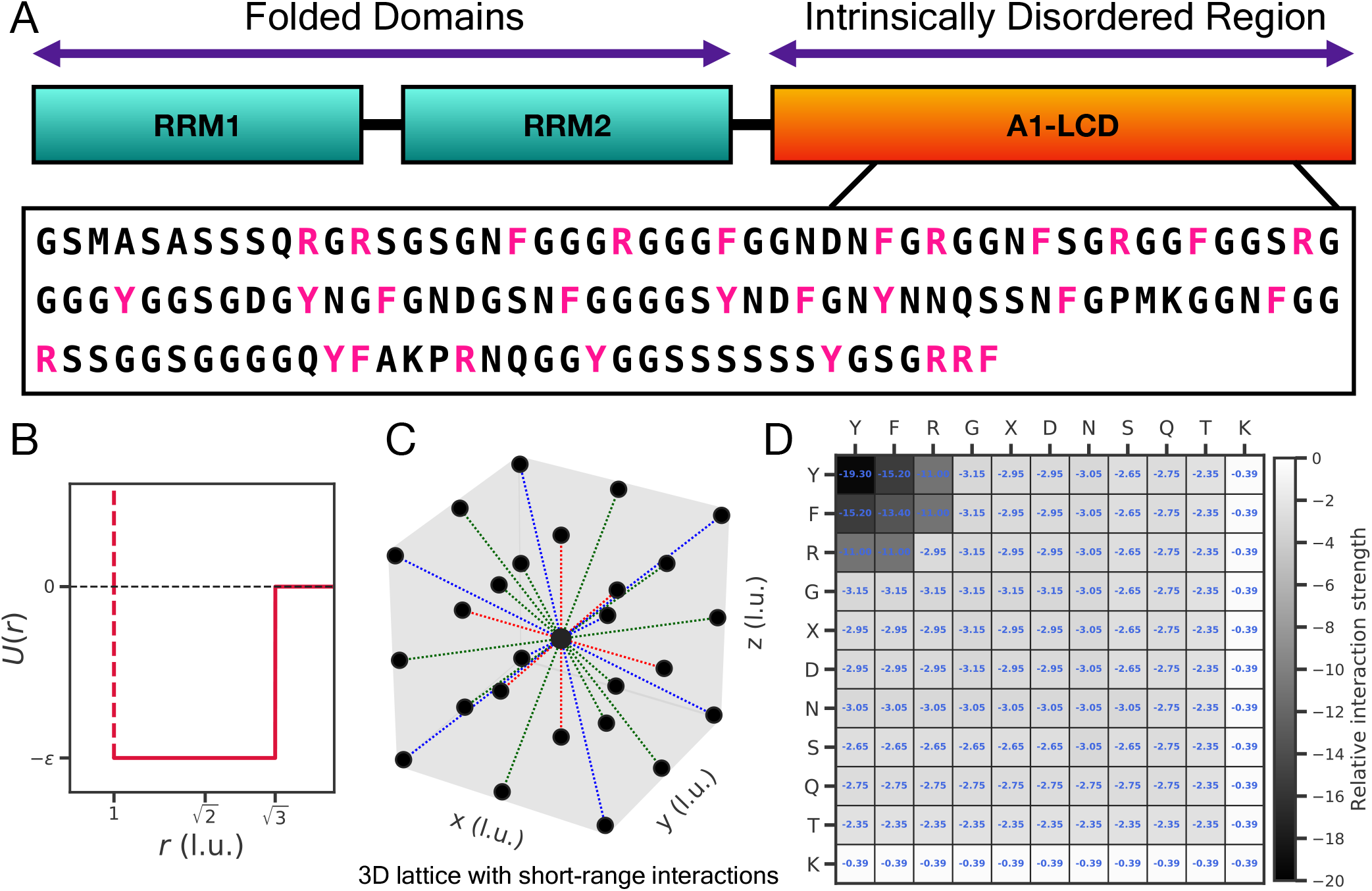
Details of the coarse-grained lattice-based Monte-Carlo simulations. (A) Architecture of the hnRNP-A1 protein and amino acid sequence of A1-LCD. hnRNP-A1 comprises the folded RNA Recognition Motifs (RRMs) and the PLCD (Prion-Like Low Complexity Domain), which is designated as A1-LCD. The amino acid sequence of A1-LCD, which has 137 residues, is also shown in the box. Sticker residues in the A1-LCD sequence were identified in previous studies (56, 60), and these are tyrosine (Y), phenylalanine (F), and arginine (R), which are colored in magenta to delineate them from the rest of the residues (colored in black). (B) Pairwise energy of interaction shown as a function of the distance between any two beads on the lattice (See Methods section). (C) Schematic of a cubic lattice with short-range interactions where each bead has 26 possible neighbors at distances of 1 l.u. (colored by red dotted lines), 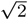 l.u. (colored by green dotted lines), or 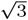 l.u. respectively (colored by blue dotted lines). (D) Matrix showing the pairwise interaction strengths used in the LaSSI computational model. “X” is used to indicate any amino acid for which a specific interaction is not defined. The values for the relative interaction strengths for each residue pair are shown inside the squares of the matrix grid. The energies are unitless. Moves are accepted or rejected by referencing to thermal energy *k*_*B*_ *T*_*s*_, where *k*_*B*_ = 1 and *T*_*s*_ is the simulation temperature.

Coarse-grained simulations have been used to map binodals for an assortment of systems (17, 20, 44, 51–57). These off-lattice molecular dynamics (MD) simulations and on-lattice Monte Carlo (MC) simulations typically include *O*(10^2^) molecules per simulation cell. In the off-lattice MD simulations, the computed dilute and dense phase arms of the binodal are connected by a numerical fitting procedure (58). The system is assumed to belong to the same universality class as the 3-dimensional Ising model. The binodal is then fit to the scaling of the order parameter prescribed by the Ising model, and the critical temperature is estimated through the fitting procedure. Here, we assess the mapping of critical points using the approach established in the literature, and then examine the accuracy of this method by performing large-scale simulations. We find it imperative to deploy rigorous finite size scaling based on analysis of Binder cumulants (59). This analysis yields accurate estimates of the critical point. We use this approach to map the full binodal of A1-LCD and demonstrate how the percolation and overlap lines intersect the coexistence curve (binodal). We identify three distinct regimes, which are characterized by distinct, regime-specific cluster-size distributions in the coexisting dilute phases. We also estimate the theta temperature as the point where the two-body interaction coefficient (*B*_2_) for a pair of chains equals zero. We find that the temperature where this criterion is met is very different from the apparent theta temperature inferred by analysis of the scaling of mean segmental distances *R*(*s*) as a function of *s*, the length of a segment.

## Methods

Our coarse-grained lattice-based Monte Carlo (MC) simulations (61–63) employ the LaSSI simulation engine (46, 56), which is based on a generalization of the bond fluctuation model (64, 65). A single A1-LCD molecule comprises 137 beads, one for each of the residues, and each residue is a single bead on a lattice. For the inter-residue potentials (Fig. 1B, Fig. 1D), we used a short-range potential function that was parameterized using a Gaussian Process Bayesian Optimization procedure (56, 66). The parameters are transferable unto other PLCDs but not to other systems (56, 57). To map the critical region, we used finite-size scaling that relies on the quantification of density fluctuations via analysis of Binder cumulants (59, 67). We used a nine-step protocol, which is followed sequentially to arrive at a complete characterization of the binodal of A1-LCD (Fig. 2). The key considerations are as follows: Obtaining converged estimates of mean densities, density fluctuations, and higher-order moments, which are imperative for mapping critical regions and percolation lines, requires that the simulation cells encompass at least ∼6 × 10^3^ A1-LCD molecules. The box sizes need to be large because of the high volume fractions near critical points, and the boxes need to be cubic with periodic boundaries to avoid confinement or symmetry-breaking artifacts. As a result, the number of MC steps required to obtain converged estimates is *𝒪* (10^11^). Based on these considerations, the simulation system includes 10^4^ A1-LCD molecules for each simulation temperature *T*_*s*_. These simulation temperatures *T*_*s*_ are converted to Kelvin by multiplying them by a factor of 5.6 (56) (see Section 2.2). For every value of *T*_*s*_, we performed at least three independent MC simulations with different random seeds. Each simulation comprises 6 × 10^11^ MC steps. In the currency of MD simulations, 6 × 10^11^ MC steps would translate to simulation times of ∼3 ms assuming an integration time step of 5 fs. Additional simulations of equivalent length (for more temperatures in the critical region) were performed when the Binder cumulant analysis needed to be deployed. For computing binodals as a function of temperature, we used cubic boxes that were 240 lattice units (l.u.) to a side. In Å, this would translate to ∼ 960 Å. For the finite-size scaling analysis, we quantified Binder cumulants using sub-boxes whose dimensions span the range of *L* ≈ 50 − 200 l.u.

**Fig. 2.**
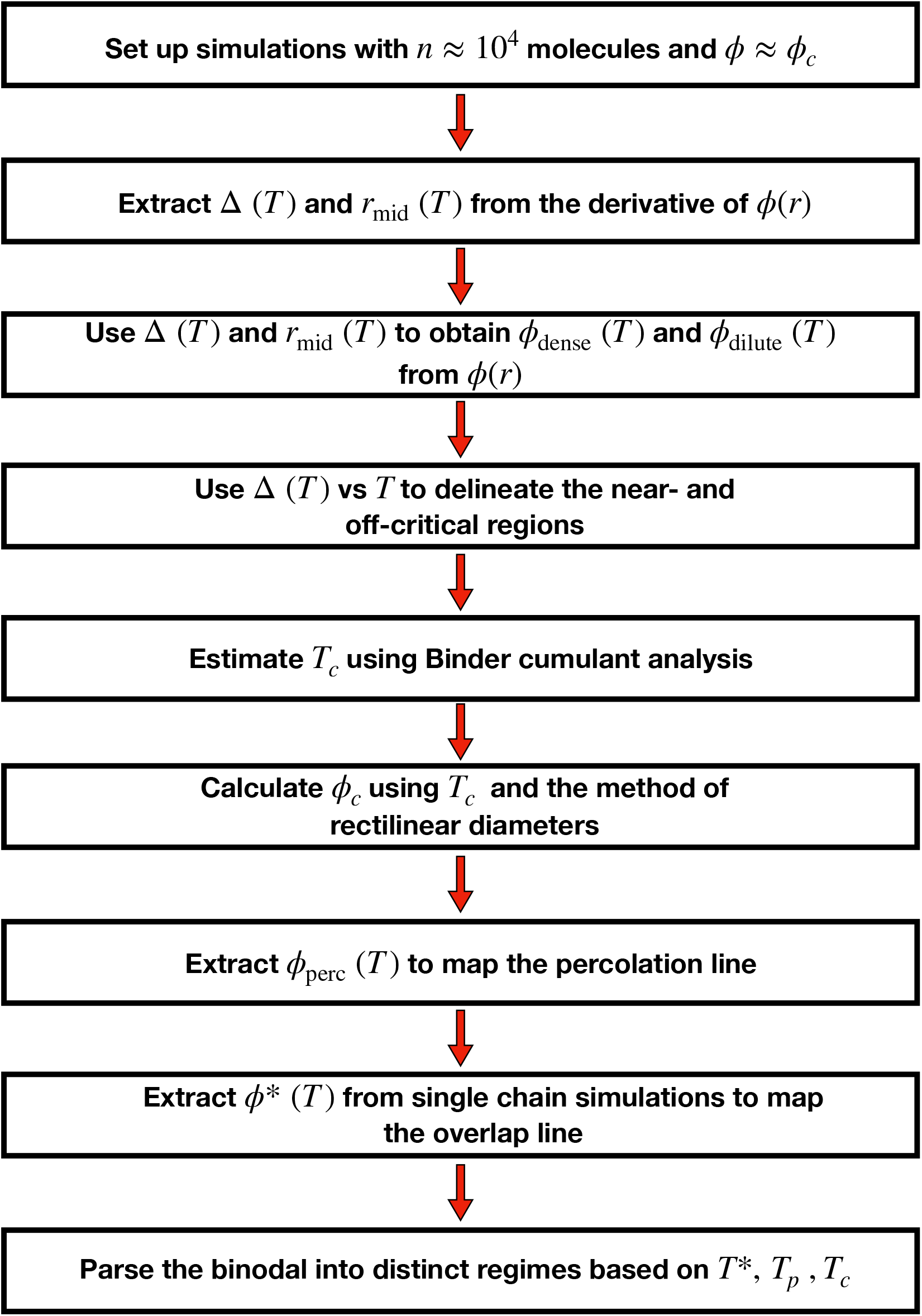
Flowchart showing the nine-step simulation and analysis protocol to map the full binodal for WT A1-LCD. Details of the protocol are provided in the Methods section of the main text.

### LaSSI simulations

The lattice-based MC simulations were performed following the approach of Farag et al., (56, 57). Beads occupy positions on a cubic lattice, and the energy of interaction is −*ε*_*ij*_ (Fig. 1B, Fig. 1C) for beads of type *i* and *j* if the distance between the beads on the lattice is 1, 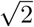, or 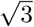 1.u. apart. Otherwise, the interaction energy is zero. The values of −*ε*_*ij*_ were those of Farag et al., (56) (Fig. 1D). The energies were parameterized to reproduce the sequence specificity of measured radii of gyration (60) and the parameterization scheme was based on the Gaussian Process Bayesian Optimization approach of Ruff et al. (66). We analyzed the results from two sets of simulations, one that includes 200 A1-LCD molecules in a cubic box (*L*_sim_ = 120 l.u.) and the other that includes 10^4^ molecules (*L*_sim_ = 240 l.u.). LaSSI simulations to study the phase behaviors were performed for 46 ≤ *T*_*s*_ *≤* 64 (257.60 K ≤ *T ≤* 358.40 K, where *T* is the temperature in Kelvin).

### System size considerations for the LaSSI simulations

To discern the optimal box size and numbers of A1-LCD molecules (*n*) that are needed to map critical regions, we first estimated the critical volume fraction (*ϕ*_*c*_) by fitting the computed binodal from the simulations with 200 molecules. For this, we used a modified Flory-Huggins model (68). The estimated value of the critical volume fraction was found to be *ϕ*_*c*_ ≈ 0.1. The contour length of each A1-LCD molecule is 137 l.u. Therefore, to achieve the estimated volume fraction using 200 molecules when the system is quenched to be just below the apparent critical point, the dimensions of the cubic box would have to be *L*_sim_ ≈ 65 l.u. to a side. However, this is less than half the chain contour length. This becomes a problem because the simulation volume of a cubic box can only accommodate density fluctuations that are smaller than the contour length of a single chain. Additionally, as *T* → *T*_*c*_, the correlation length *ξ*_*c*_ becomes comparable to the box size *L*_sim_. Therefore, a box comprising 200 molecules with *L*_sim_ ≈ 65 l.u. poses two challenges: First, the estimates of density fluctuations will be large due to finite-size effects resulting from the use of small numbers of molecules to mimic a macroscopic system. Second, the long-wavelength density fluctuations will be suppressed by the small box size that is necessitated by the use of only 200 molecules. In contrast, for 10^4^ molecules, the box length will need to increase to *L*_sim_ ≈ 239 l.u. to achieve a volume fraction of ≈ *ϕ* 0.1. This box size accommodates many independent correlation volumes thereby enabling the sampling of a broader spectrum of density fluctuations.

To estimate the appropriate numbers of molecules and box sizes, we used the following considerations. For fixed *ϕ* ≈ *ϕ*_*c*_, the adequate box length will scale as *L*_sim_ ∝ *n*^1*/*3^, where *n* is the number of A1-LCD molecules. Since 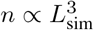, to determine the minimum number of molecules that would be required to map phase behaviors close to criticality, the box length *L*_sim, target_ at *ϕ* ≈ *ϕ*_*c*_ should be between 1.5 × *N* and 2 × *N*, where *N* = 137 is the contour length for A1-LCD. This gives

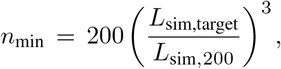

where *L*_sim,200_ = 65 l.u. and *L*_sim,target_ = 1.5 × 137 l.u. or *L*_sim,target_ = 2 × 137 l.u. Substituting these values, we obtain

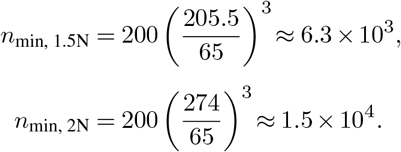

Simulations with *n* ≈ 10^4^ lie within the prescribed range, and this should be adequate to map the phase behavior in the vicinity of the critical point. In our simulations, we used a cubic box of size *L*_sim_ = 240 l.u. for a system of 10^4^ A1-LCD molecules.

### Conversion from simulation to experimental units

Scaling factors are required to convert the computed coexistence curves from simulation temperatures into degree Kelvin and volume fractions into molar units. For the latter, we followed Wei et al. (38) and Farag et al. (56). A volume fraction of 1.0 corresponds to a mass concentration of 1310 mg/ml. Following Farag et al. (56), we used a multiplicative factor of 5.6 to convert the simulation temperature in LaSSI (which is written in units of *k*_*B*_ = 1) to degrees Kelvin. Farag et al. derived this factor as follows: Binodals extracted from LaSSI simulations were converted into mass concentrations along the abscissa, and the ordinate was multiplied by a conversion factor that maximized the overlap with the measured binodals for A1-LCD (17, 56). This conversion, which was derived for A1-LCD, was then applied without further modification to more than 30 different sequences derived from the A1-LCD parent (56, 60) and to the LCD of FUS (57). A fixed density scaling factor (0.6) was used to convert volume fraction from lattice units to volume fraction as measured experimentally or, equivalently, from simulation concentrations in (M) to experimental concentrations measured in (M) (17, 56). Lengths in lattice units were converted to real units using a lattice spacing of 4 Å per lattice unit. This spacing was chosen because it corresponds to the average distance between adjacent C_*α*_ atoms (3.8 − 4.0 Å) (69, 70). Consequently, one lattice site represents a single residue both in volume and in bond length, yielding physically realistic chain dimensions. Together, these conversions enable direct comparison between lattice simulations and experimental data.

### Protocol for accurately mapping the critical regime and estimating critical parameters

**Step 1** involves setting up simulations of the appropriate size (box size and numbers of molecules) and length (number of MC steps). **Step 2** is the extraction of the interfacial width Δ(*T* ) and the midpoint of the interface *r*_mid_ from the radial density profiles. **Step 3** uses the *T* -dependent radial density profiles and the estimated *T* -dependent values of Δ(*T* ) and *r*_mid_ to extract *ϕ*_dense_(*T* ) and *ϕ*_dilute_(*T* ) using the Fisk-Widom function (71). **Step 4** delineates the critical and off-critical regimes by analyzing the variation of Δ(*T* ) with *T*, quantifying *T*_crossover,Δ_, and designating the region above *T*_crossover,Δ_ as the critical region. **Step 5** deploys the analysis of Binder cumulants, which requires considerable additional sampling for temperatures closer to the critical region, to estimate *T*_*c*_. This knowledge can be used to estimate *ϕ*_*c*_ directly using the full binodal. As an additional route to estimating *ϕ*_*c*_, **Step 6** uses the known *T*_*c*_ to calculate *ϕ*_*c*_ via the method of rectilinear diameters. **Step 7** deploys sampling at high densities to extract the temperature-dependent percolation thresholds, the locus of which constitutes the percolation line. **Step 8** estimates the overlap line by performing single-chain simulations at different temperatures to estimate *ϕ*^*^(*T* ). **Step 9** quantifies the percolation and overlap lines, and estimates the points where these lines intersect the binodal. All steps of this protocol are shown in the flowchart in Fig. 2.

### Estimation of the coexistence concentrations and the interface width

To determine the interfacial width Δ and the midpoint of the interface *r*_mid_, we analyzed the derivatives of the radial density profile *ϕ*(*r*). The profile of *dϕ*(*r*)*/dr* exhibits a peak centered at the location of the interface. We fit this peak to a Gaussian function (72),

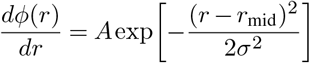

where *r*_mid_ is the midpoint of the interface and *σ* is the width of the Gaussian. The full width at half maximum (FWHM) of the fitted Gaussian is taken to be the interfacial width, 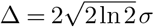. The dilute- and dense-phase concentrations used to construct the binodal are obtained by fitting the radial density profile *ϕ*(*r*) to the Fisk–Widom function (71), which interpolates smoothly between the two bulk concentrations and accounts for the interfacial width. The functional form is

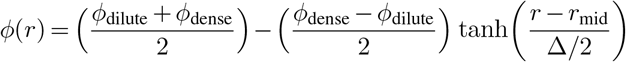

where *ϕ*_dilute_ and *ϕ*_dense_ denote the dilute- and dense-phase concentrations, respectively. The midpoint *r*_mid_ and interfacial width Δ are those extracted from the derivative fit. The coexistence concentrations *ϕ*_dilute_ and *ϕ*_dense_ reported for the binodal are obtained by averaging over three independent replicate simulations for each temperature. As an important aside, we note that Farag et al., (56) used the Fisk-Widom equation and estimated four parameters, *ϕ*_dilute_, *ϕ*_dense_, Δ, and *r*_mid_, simultaneously from fits to the radial density profiles. This approach yielded confounding observations regarding the chain-length (*N* ) dependence for the scaling of Δ with *T* for homopolymers. Specifically, Farag et al. observed an *N* - dependence for Δ(*T* ) for *T < T*_crossover,Δ_, and a convergence above this temperature. In hindsight, the correct behavior should be the opposite of what was observed and reported as being confounding (56). Closer scrutiny revealed that the erroneous observation stems from the extraction of four parameters from a single fit. Such a fit is weighted toward the high density of points in the plateau regions, which define the densities of the coexisting phases. To eliminate this error, we adopted the two-step approach introduced by Chauhan et al., (72). We first used the derivatives of the temperature-specific radial density profiles *ϕ*(*r*) to estimate Δ(*T* ) and *r*_mid_(*T* ) (see Fig. S1 in *SI Appendix*). Next, we use the derived values of Δ(*T* ) and *r*_mid_(*T* ) in the Fisk-Widom function to estimate *ϕ*_dilute_ and *ϕ*_dense_.

### Binder cumulants

We used the Binder cumulant analysis to estimate *T*_*c*_. This involves sampling a large number of sub-boxes of size *L* within the full simulation box of size *L*_sim_ and computing the density distribution and moments of the distribution, which include the mean density, the variance, and kurtosis within each sub-box. The variance and kurtosis are then used to evaluate the Binder cumulants. Analysis of the *L*-dependent cumulants as a function of *T* leads to an estimate of *T*_*c*_ as the temperature of intersection of the different cumulant curves. The order parameter of interest at a given temperature *T* is the deviation of the density inside a cubic sub-box of length *L* from its mean value:

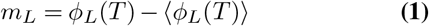

We define the second and fourth central moments of the sub-box order parameter as 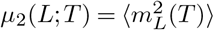 and 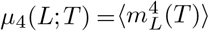, respectively. The dimensionless Binder cumulant is then

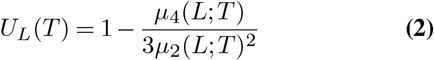

or equivalently, *U*_*L*_ = 1 − *κ/*3, with *κ* denoting the kurtosis of *p*(*m*_*L*_). In the one-phase regime above *T*_*c*_, *p*(*m*_*L*_) should be Gaussian and unimodal, corresponding to *U*_*L*_ → 0. In the two-phase regime, *p*(*m*_*L*_) becomes bimodal, and *U*_*L*_ → 2*/*3 (59, 67, 73). To choose the optimal range of sub-box sizes, we used: *R*_*ee*_ *< L* ≲ 0.8 *L*_sim_. Here *L*_sim_ is the length of the full simulation box (*L*_sim_ = 240 l.u. for a system of 10^4^ A1-LCD molecules). This choice of upper and lower bounds for *L* ensures that each sub-box is larger than the mean end-to-end distance of a single polymer (*R*_*ee*_), while remaining small enough relative to *L*_sim_ to provide sufficient statistical sampling of density fluctuations. The upper limit of *R*_*ee*_ corresponding to the highest temperature in our simulations is ≈ 17 l.u. as shown in Fig. S2 in *SI Appendix*. Thus, the range of sub-box sizes we used for the Binder cumulant analysis (*L* ≈ 50 − 200 l.u.) fall within acceptable limits.

### Sampling of sub-boxes and calculation of sub-box densities

10^4^ cubic sub-boxes of size *L* were placed uniformly at random locations within the simulation box. The coordinates of the origin of each sub-box (*x*_0_, *y*_0_, *z*_0_) were drawn independently from a uniform distribution: (*x*_0_, *y*_0_, *z*_0_) ∼ 𝒰 (0, *L*_sim_ − *L*), ensuring that all sub-boxes remained entirely within the simulation boundaries. Random uniform draws produce overlapping sub-boxes, which provide an unbiased and statistically dense sampling. The simulation coordinates were discretized onto a cubic lattice of size 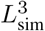, producing a binary occupancy grid *ϕ*(**r**) that equals 1 for occupied sites and 0 otherwise. The list of bead coordinates was ordered sequentially by chain. For each randomly placed sub-box *Vi* of volume *L*^3^, the local density was calculated as 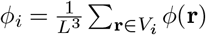, corresponding to the fraction of occupied lattice sites within that region. The sub-box densities were used to construct the probability distribution *p*_*L*_(ϕ) and to compute the second and fourth-order moments of the order parameter.

### Pairwise Binder Cumulant crossings to determine T_c_

Binder cumulant curves *U*_*L*_(*T* ) from different sub-box sizes *L* intersect near the fixed-point value *U* ^*^ (59). To estimate the critical temperature *T*_c_, we employed the standard Binder cumulant crossing analysis across sub-boxes of size *L*. For each sub-box of size *L*, the discrete values of *U*_*L*_(*T* ) obtained from simulations were interpolated using a monotonic piecewise cubic Hermite interpolant, yielding continuous *U*_*L*_(*T* ). For every pair of sub-box sizes (*L*_1_, *L*_2_) 160 l.u., we computed the difference 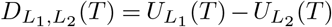, and located all temperatures *T* satisfying 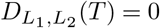. For this, we used Brent’s root-finding method (74) within the overlap of their temperature ranges. The resulting set of pairwise crossing temperatures 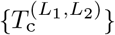 can be treated as a distribution of estimates for the critical temperature. The median of all filtered pairwise crossings thus provides a robust estimate of *T*_c_ with well-defined uncertainty bounds (the upper and lower values of this set).

### Percolation threshold as a function of temperature

To determine the percolation threshold *ϕ*_perc_ at a given temperature, we compute the percolation order parameter defined as the fraction of chains belonging to the largest connected cluster (lc), following the approach used by Choi et al., (46),

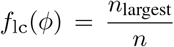

Here, *n* = 10^4^ is the total number of A1-LCD molecules in the system. The resulting *f*_lc_(*ϕ*) versus *ϕ* curve exhibits a sigmoidal shape, characteristic of a percolation transition. Accordingly, we fit the data to a logistic function of the form

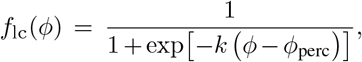

where *k* is a parameter describing the steepness of the transition and *ϕ*_perc_ denotes the percolation threshold. The value of *ϕ*_perc_ is identified with the inflection point of the fitted logistic curve, corresponding to *f*_lc_(*ϕ*) = 0.5. We deploy this computation of *ϕ*_perc_ only for state points corresponding to temperatures that lie outside the two-phase region of the binodal for WT A1-LCD (see Fig. S4 in *SI Appendix*).

### Calculation of the overlap volume fraction from single chain simulations

The overlap volume fraction *ϕ*^*^, which is a function of temperature, is obtained from single-chain simulations by first computing the root-mean-square end-to-end distance (see Fig. S2 and Fig. S5 in *SI Appendix*),

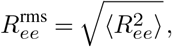

and then estimating the coil volume as 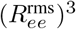. The overlap volume fraction is defined as

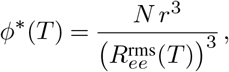

where *N* = 137 is the number of residues in a chain and *r* = 0.5 l.u., the radius of individual residues/beads.

### Defining clusters based on connectivity of stickers

Clustering of A1-LCD chains is quantified using a sticker-mediated connectivity criterion applied to individual simulation snapshots. “Sticker” residues (17, 29, 60, 75) are defined as tyrosine (Y), phenylalanine (F), and arginine (R), as these residues are known to be primary drivers of intermolecular interactions in PLCDs (17, 60). The choice of these residues as stickers is supported by the hierarchy of interaction strengths encoded in the simulation energy matrix (see Fig. 1D), wherein Y–Y, F–F, Y–F, and R–Y, R–F interactions constitute the most favorable non-bonded contacts. Accordingly, only beads corresponding to Y, F, and R residues are considered when constructing intermolecular contacts. For each snapshot, sticker bead coordinates are wrapped under periodic boundary conditions and inserted into a lattice-based spatial lookup table. Two chains are considered connected if at least one sticker bead from one chain occupies a nearest-neighbor lattice site (including axial and diagonal neighbors, corresponding to distances of 1, 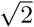, or 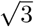 l.u.) of a sticker bead from another chain. This procedure generates a chain-level adjacency graph in which nodes represent chains and edges represent at least one inter-chain sticker contact (56, 76, 77). Connected components of this graph are identified using a depth-first search (DFS) algorithm, and each connected component is defined as a cluster. Note that even at the highest concentrations, less than 10% of the lattice volume is taken up by chain molecules. Additionally, the bulk A1-LCD concentrations are identical across all temperatures, implying that the cluster-size distributions that we extract from the simulations are emergent properties of the phase behaviors and not a trivial consequence of including 10^4^ molecules in the simulations.

### Calculation of the scaling exponent *ν*(*T* ) from the ensemble-averaged segmental distances *R*(*s*)

For each temperature *T*, the ensemble-averaged internal scaling profile *R*(*s*) was first obtained by averaging over equilibrated simulation snapshots and across three independent replicates. The scaling exponent *ν* was then extracted from this temperature-specific profile using a finite-difference formulation in logarithmic space. Specifically, for segment pairs corresponding to contour separations *s*_*l*_ and *s*_*u*_, we computed

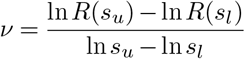

following the approach described in (78). This expression estimates the local slope of ln *R*(*s*) versus ln *s*, thereby yielding an effective scaling exponent without imposing a global regression. The analysis was restricted to intermediate segmental separations satisfying 40 ≤ *s*_*l*_, *s*_*u*_ *≤* 120. The lower bound excludes locally stiff segments that do not obey asymptotic scaling behavior, whereas the upper bound avoids artifacts arising from finite chain length and dangling chain ends. Only segment pairs satisfying *R*(*s*_*u*_) *> R*(*s*_*l*_) were retained to ensure physically meaningful positive slopes. This procedure yields a distribution of *ν* values at each temperature, from which the mean and standard deviation were computed. The scaling analysis was performed only for temperatures above the “globule” regime where the internal scaling profiles become effectively flat. At lower temperatures, strong chain collapse results in nearly *s*-independent *R*(*s*) profiles, rendering the logarithmic derivative ill-defined and thus preventing the meaningful extraction of a scaling exponent. We therefore, restrict the scaling analysis to all temperatures *T >* 140 K.

### Computing the temperature dependence of the two– body interaction coefficient *B*_2_

To determine the temperature dependence of the two-body inter-chain interaction coefficient *B*_2_ for A1-LCD, we performed umbrella sampling simulations of a pair of chains across a range of temperatures *T* . The reaction coordinate was defined as the separation between the centers-of-mass of the two chains. Harmonic biasing potentials were applied at multiple restraint distances to ensure sufficient overlap between adjacent windows (see Fig. S6 in *SI Appendix*). The Weighted Histogram Analysis Method (WHAM) (79–81) was then used to combine biased sampling data from multiple overlapping umbrella windows to self-consistently reconstruct the unbiased potential of mean force *W* (*r*) for each temperature. The Mayer *f* -function is defined as (82, 83):

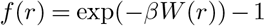

where 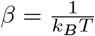. The temperature-dependent two-body interaction coefficient is proportional to the excluded volume v_ex_ ∝ *B*_2_:

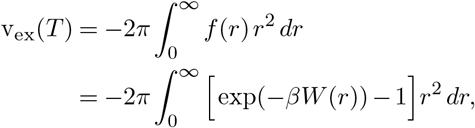

where the upper limit of integration is chosen such that *W* (*r*) → 0. Repeating this procedure at different temperatures yields v_ex_(*T* ) ∝ *B*_2_(*T* ), thus providing a quantitative measure of how the effective pairwise attractions between A1-LCD chains evolve with temperature. We compute v_ex_(*T* ) and then estimate *B*2′ (*T* ) as the ratio of *B*_2_(*T* ) at a temperature *T* to the magnitude of *B*_2_ corresponding to the highest temperature in our simulations.

## Results

### Assessing finite-size artifacts

It has become common practice to assume that the order parameter defined by Δ*ϕ* = (*ϕ*_dense_ − *ϕ*_dilute_) scales as 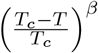, with *β* set to be ≈ 0.33, being the exponent that is expected for the 3D Ising model. Here, *ϕ*_dense_ is the volume fraction of the dense phase, *ϕ*_dilute_ is the volume fraction of the dilute phase, *T* is the simulation temperature, and *T*_*c*_ is the critical temperature. The procedure that is typically followed is to use the law of rectilinear diameters and the assumption of the scaling of the order parameter to be congruent with the 3D Ising model, thereby fixing *β*, and using a regression analysis to estimate *T*_*c*_, *ϕ*_*c*_, and a pre-factor *A*_0_ (20, 51, 53–55, 84, 85). Note that the scaling of the order parameter is only valid in the vicinity of the critical point (86). Accordingly, some authors exercise caution in the use of the regression analysis (87), although this will be affected by the paucity of points that are sampled in the vicinity of the critical point from simulations of finite-sized systems.

To assess whether the vicinity of the critical regime is actually being sampled in simulations of finite-sized systems, we examined the *T* -dependent radial density profiles for a system of 200 A1-LCD molecules (Fig. 3A). A defining feature of the critical regime is that the concentration of the dilute phase should approach that of the dense phase. Concentrations extracted from these profiles using the Fisk-Widom function show that at all temperatures sampled, the dilute and dense phase concentrations differ by over an order of magnitude (circles in Fig. 3B, Fig. 3C). This suggests a lack of data points around the critical point. Additionally, as described in the Methods section, sampling the critical regime for a simulation with 200 molecules of A1-LCD would require a box size of *L*_sim_ ≈ 65 l.u. which imposes finite-size artifacts.

**Fig. 3.**
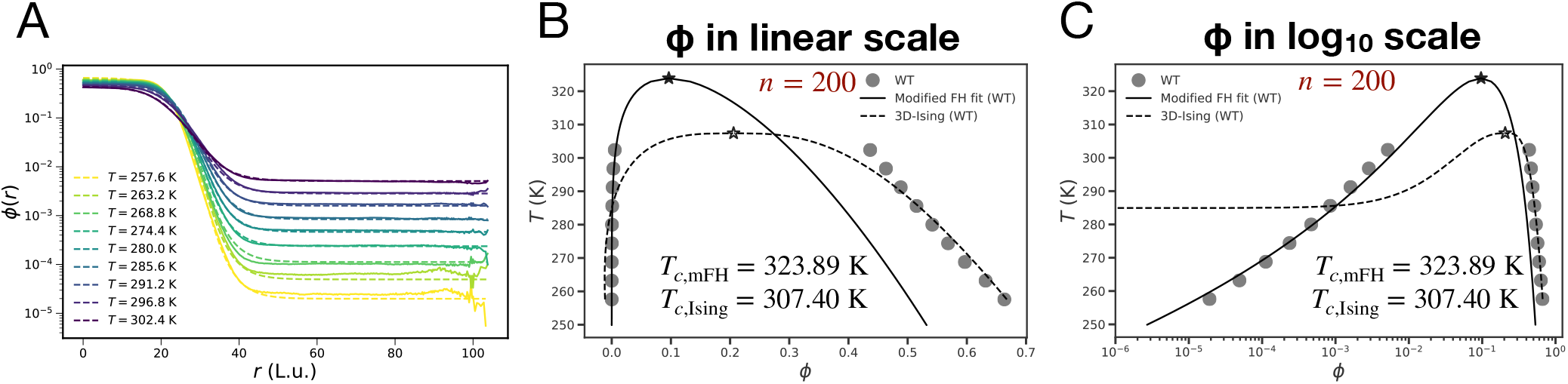
Finite-sized systems pose limitations in mapping the full binodal. The results are shown here for A1-LCD. (A) *T* -dependent radial density profiles *ϕ*_*T*_ (*r*) for the system containing 200 molecules of WT A1-LCD. Different profiles were obtained for different temperatures *T* (in Kelvin). For each temperature, the computed radial density profile was used to extract the dense and dilute coexistence concentrations by fitting them to the Fisk-Widom function. (B-C) Comparative analysis of estimates of critical points and fitting binodals to the modified FH model and the 3D-Ising model. The results are shown in (B) linear scale and (C) log scale of the volume fraction *ϕ*. These results highlight the weaknesses of simulations based on finite-sized systems and the use of fits to analytical expressions without assessing the contributions from finite size artifacts.

To work around this paucity of points, the common practice is to enforce the scaling of the order parameter across the entirety of the computed binodal. The reduced temperature is defined as

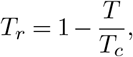

where *T*_*c*_ is the critical temperature. According to the law of rectilinear diameters, the average of the coexisting phase volume fractions varies linearly with temperature:

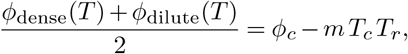

where *ϕ*_*c*_ is the critical volume fraction and *m* is the slope. Next, the difference between the dense and dilute arms is taken to scale as a power law with Ising critical exponent *β* ≈ 0.33:

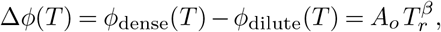

where *A*_*o*_ is a non-universal amplitude. Combining these two expressions, the binodal branches are written as

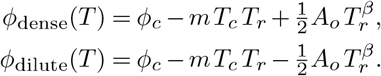

The system of equations combined with setting *β* ≈ 0.33, and knowledge of the computed values of *ϕ*_dense_(*T* ) and *ϕ*_dilute_(*T* ) across the entire binodal are used to estimate *m, T*_*c*_, *ϕ*_*c*_, and *A*_0_.

We assessed the quality of the fits obtained using the approach laid out above by utilizing the 200 chain A1-LCD system (Fig. 3B, Fig. 3C). The dilute arm is fit very poorly by imposing the scaling of the order parameter from the 3D Ising model to apply across the entire binodal. This is congruent with the fact that this scaling should only apply in the vicinity of the critical point. Next, we fit the computed binodal to a modified Flory-Huggins model (17, 60, 68). While this method generates a mean field scaling of the order parameter in the vicinity of the critical point (88), its use for estimating the location of the critical regime is appropriate. The binodal obtained by fitting the data to the modified Flory-Huggins model does a better job, especially in terms of fitting the dilute arm. The estimated *T*_*c*_ is considerably higher than what is estimated by imposing scaling of the order parameter based on the 3D Ising model. These discrepant estimates, along with the radial density profiles, suggest that the critical regime is inadequately sampled or not sampled at all in simulations based on 200 A1-LCD molecules, thus emphasizing the challenges posed by finite size artifacts (89) (Fig. 3).

### Mapping the binodal for large systems and assessing its accuracy

For the simulations that include 10^4^ A1-LCD molecules, the presence of a two-phase system in the simulation cell is evident from the temperature-specific radial density profiles *ϕ*(*r*) (Fig. 4). A hyperbolic tangent, Fisk-Widom function (56, 71) is fit to the *T* -dependent profiles for *ϕ*(*r*) to estimate *ϕ*_dense_(*T* ) and *ϕ*_dilute_(*T* ) for the system of 10^4^ A1-LCD molecules (Fig. 4). The simulations for a system with 10^4^ chains show that the envelope of the two-phase regime extends to higher temperatures. As *T* increases towards the apparent critical regime, we observe nearly equivalent dilute and dense phase volume fractions (Fig. 5A), which could not be observed in simulations of the 200-chain system. To assess the accuracy of the computed binodals, we compared them to one another and to experiments. Note that the experiments used absorbance for measuring off-critical coexistence points, and cloud point measurements to map the critical region (60). The cloud point measurements suggest that the critical region is well above *T*_*c*,Ising_ = 307.40 K, as estimated from the 200-chain simulations. However, the cloud point measurements are noisy due to large density fluctuations. Hence, the experiments do not yield an accurate measurement of *T*_*c*_, but they identify the near-critical regime. To put the comparisons between computed and measured binodals on a quantitative footing, we computed transfer free energies using 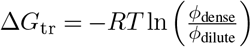 in units of kJmol^−1^ (90). In the regime where the comparisons are feasible, the computed transfer free energies from both the 200-chain and 10^4^-chain systems showed good agreement with the measurements (Fig. 5B). These comparisons give us confidence regarding the accuracy of the model used in the LaSSI simulations.

**Fig. 4.**
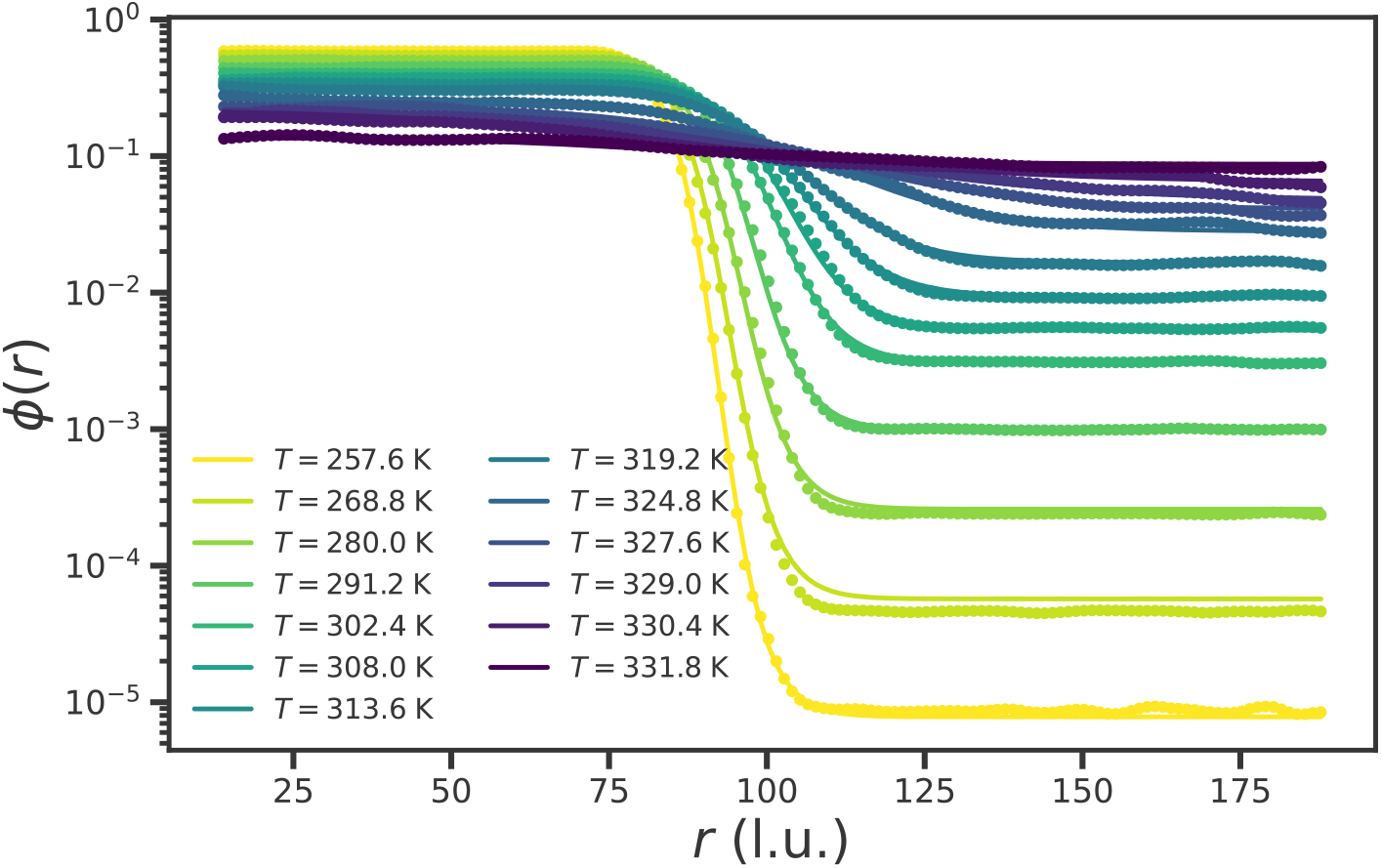
T -dependent radial density profiles for a system with 10^4^ molecules of A1-LCD. Radial density profiles *ϕ*(*r*) as a function of *r* (in lattice units) at different temperatures for the 10^4^ chain system, which are then fit using the Fisk-Widom function to estimate the coexisting dense and dilute phase concentrations (*ϕ*_dense_ and *ϕ*_dilute_).

**Fig. 5.**
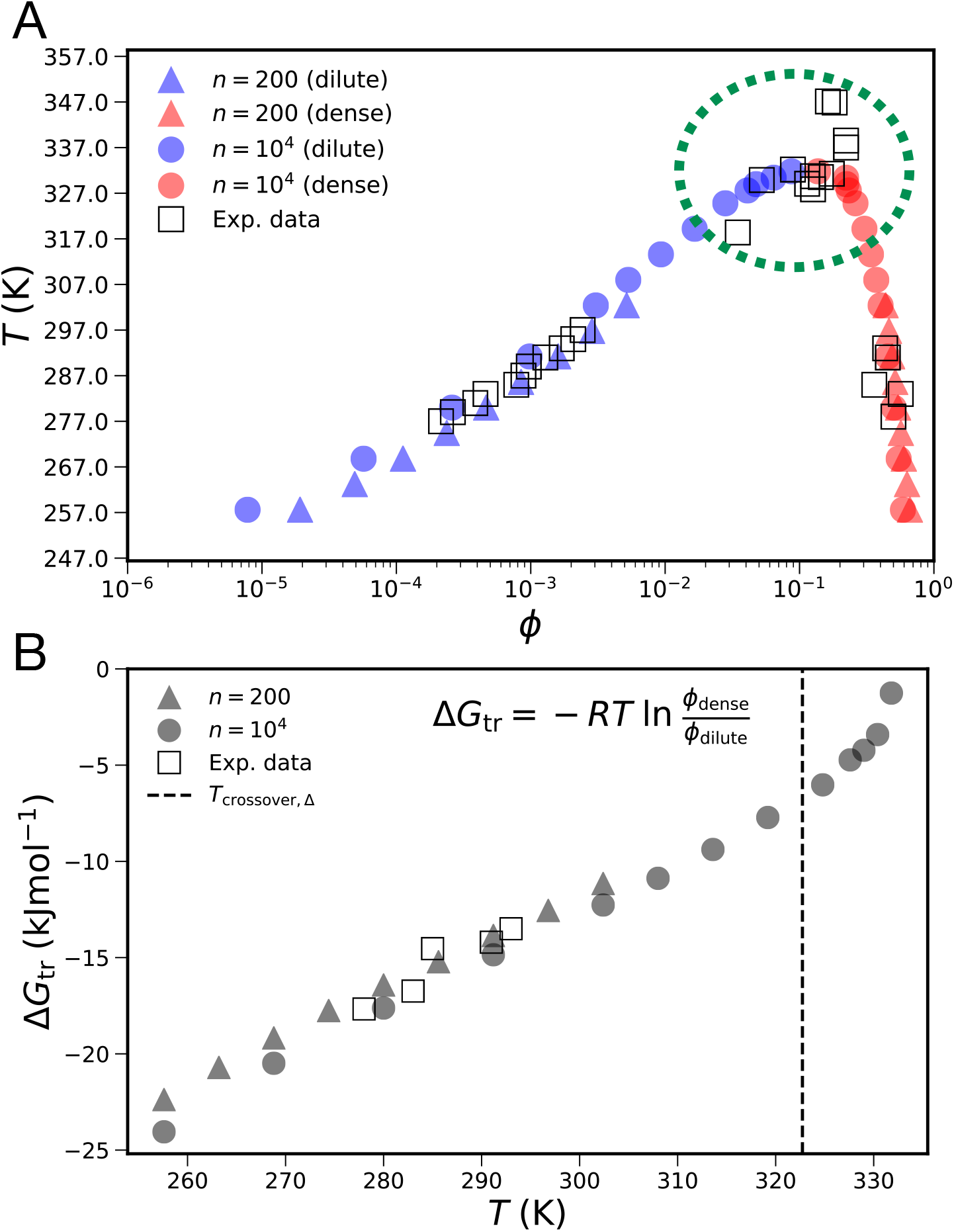
Comparisons between computed and measured binodals. (A) Binodal of A1-LCD, showing coexistence concentrations of the dilute phase (blue) and dense phase (red). Results from simulations with 10^4^ chains are shown as circles, and those from simulations with 200 chains are shown as triangles. The measured coexistence points (60) are shown as open squares. The critical regime is delineated by the green dotted oval. In this regime, the experiments reveal large density fluctuations. In the simulations, the larger system size enables accurate mapping of the critical region, and the binodal is well-rounded near the apparent critical point. Conversely, simulations based on the 200-chain system cannot access the critical regime. This emphasizes the erroneous nature of the estimates of *T*_*c*_ and *ϕ*_*c*_ that we obtain from binodals for the 200-chain system. (B) To assess the accuracy of the binodal, we computed transfer free energies, calculated as Δ*G*_tr_ = −*RT* ln(*ϕ*_dense_ */ϕ*_dilute_ ). These are plotted as a function of temperature. Here, *R* = 8.314 Jmol^−1^ K^−1^ . Triangles correspond to data from the 200-chain simulations, circles to data from 10^4^ -chain simulations, and squares to measurements. The dashed vertical line corresponds to *T*_crossover,Δ_ ≈ 322.74 K between the mean field and critical regimes (see text and Fig. 7B).

Fitting the entire binodal for the 10^4^-chain system to the scaling of the order parameter that is based on the 3D Ising model yields a higher estimate of *T*_*c*_, when compared to the 200-chain system. However, the reproduction of the binodal is very poor (Fig. 6) and this is expected because the scaling is invalid away from the critical regime. Indeed, when we restrict the analysis to the near-critical regime, i.e., to temperatures above *T*_crossover,Δ_ (see Fig. 7B), the estimated *T*_*c*_ does not change much. Even here, the symmetry expected of the order parameter is not present in the actual data, and hence the fit to the binodal in the vicinity of the critical point remains poor (Fig. 6B). As for the modified Flory-Huggins model, the fits, in general, look better than what we obtain based on the 3D Ising model. However, the value of *β* will always be 0.5, and this goes against expectations of behaviors in the vicinity of the critical point (82, 88).

**Fig. 6.**
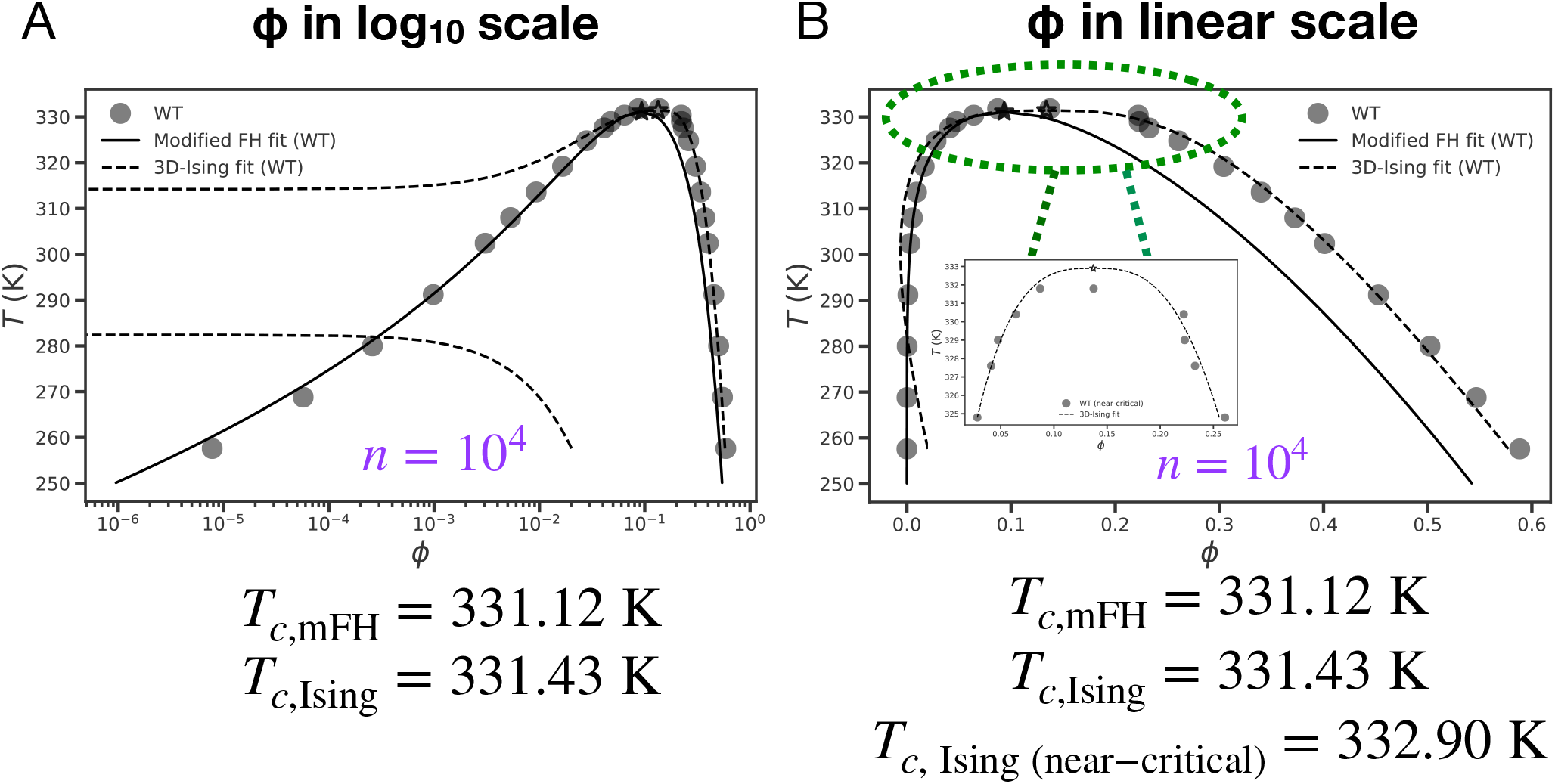
Illustration of the error of using a single model to fit the entire binodal. Comparative analysis of estimates of critical points based on two different approaches. Fits performed using the modified FH model and the 3D-Ising model to the entirety of the binodal for a system comprising 10^4^ molecules of WT A1-LCD. The results are shown in both (A) log scale and (B) linear scale of the volume fraction *ϕ*. The inset in (B) shows results obtained by fitting the 3D-Ising model to points in the critical regime, i.e., *T > T*_crossover,Δ_ (see Fig. 7B). Even though the analysis has access to concentrations in the critical regime, the fit to the dilute arm using the 3D-Ising model approach is poor.

**Fig. 7.**
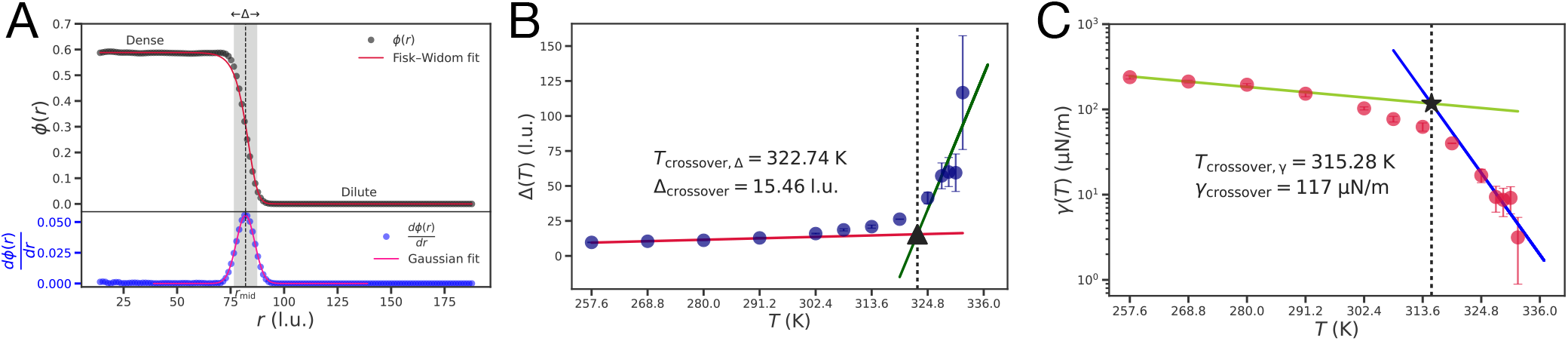
Demarcation of the off-critical (mean field) and near-critical regimes. (A) For a given temperature, the derivative of the radial density profile *ϕ*(*r*) is fitted to a Gaussian function, from which the interfacial width Δ is extracted as the full-width-at-half-maximum (FWHM). The midpoint of the interface *r*_mid_ is the mode of the Gaussian. We insert the computed values of Δ and *r*_mid_ into a hyperbolic tangent Fisk-Widom function (56, 71). This function is then fit to the *ϕ*(*r*) profile to estimate the coexisting dense and dilute-phase concentrations (*ϕ*_dense_ and *ϕ*_dilute_). (B) Temperature dependence of Δ(*T* ) showing a crossover from the off-critical to the near-critical regime at *T*_crossover,Δ_ ≈ 322.74 K, with a corresponding interface width Δ_crossover_ ≈ 15.5 lattice units. (C) A similar crossover is observed for the interfacial energy per unit area *γ*(*T* ), with 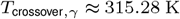 and *γ*_crossover_ ≈ 117 *µ*N*/*m.

In summary, the preceding analysis establishes why estimation of the critical parameters and / or characterizations of near-critical behaviors (44) are likely to be problematic if one uses approaches that are in vogue in the computational literature. Of course, as crude estimates of *T*_*c*_ and for assessing how this parameter varies with changes to sequence, the approach of regression is likely to be useful. For this exercise, we recommend the use of fits to the modified Flory-Huggins model (17), as opposed to the use of regression based on assuming scaling of the order parameter based on the 3D Ising model (54, 55). At least with the modified Flory-Huggins model, one gets a more accurate estimation of the transfer free energy, and this provides greater confidence with regard to comparative assessments of sequence-specific effects on phase behaviors.

### Delineating the critical and off-critical regimes

At a given temperature *T*, coexisting dense and dilute phases are delineated by an interface of width Δ. In the critical regime, the system is defined by large density fluctuations, and Δ will diverge as *T* → *T*_*c*_ (44, 91). The derivative of *ϕ*(*r*) with respect to *r* (Fig. 7A) helps delineate the location of the interface. We inserted the computed values of Δ(*T* ) and *r*_mid_(*T* ) into a hyperbolic tangent, Fisk-Widom function (56, 71). This function is then fit to the *T* -dependent profiles for *ϕ*(*r*) to estimate *ϕ*_dense_(*T* ) and *ϕ*_dilute_(*T* ) (Fig. 4, Fig. 7A).

The width Δ(*T* ) shows the presence of two regimes as a function of temperature, with the crossover temperature being *T*_crossover,Δ_ ≈ 322.74 K (Fig. 7B). Above *T*_crossover,Δ_, Δ(*T* ) increases rapidly with *T* . This is characteristic of the divergence one expects for the width of the interface as the critical temperature *T*_*c*_ is approached (44, 91). Thus, we used *T*_crossover,Δ_ to demarcate the critical and off-critical regimes, with *T > T*_crossover,Δ_ designated as the critical regime. The interfacial tension is the free energy penalty associated with increasing the area of the interface and it is computed as 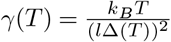 (Fig. 7C) (92). Here, *l* = 4 is a factor that converts 1 l.u. to 4 Å (see Methods section). The interfacial tension shows a sharp decrease above *T*_crossover,Δ_ and it is consistent with the expectation that *γ*(*T* ) will approach zero as *T* → *T*_*c*_ (44). The calculated values of *γ*(*T* ) are consistent with measured interfacial tension values for condensates (4, 40, 93–95) and for colloid-polymer systems (92).

### Analysis of Binder cumulants to estimate *T*_c_

To avoid the problems outlined in the preceding sections and take advantage of the accuracy afforded by large-scale simulations, we used an approach based on Binder cumulants, which rests on computing density cumulants and the use of rigorous finite-size scaling (59, 67, 89, 96–98). The Binder cumulants quantify the mean density (first moment), density fluctuations (based on the second moment), and the anisotropic nature of the fluctuations (based on the fourth moment). The finite-size scaling uses cubic sub-boxes, each of a different length *L*. Each sub-box of length *L* is used to probe the mean density and cumulants on the length scale of the sub-boxes for all temperatures. A schematic of the workflow deployed to calculate and analyze Binder cumulants is shown in Fig. 8.

**Fig. 8.**
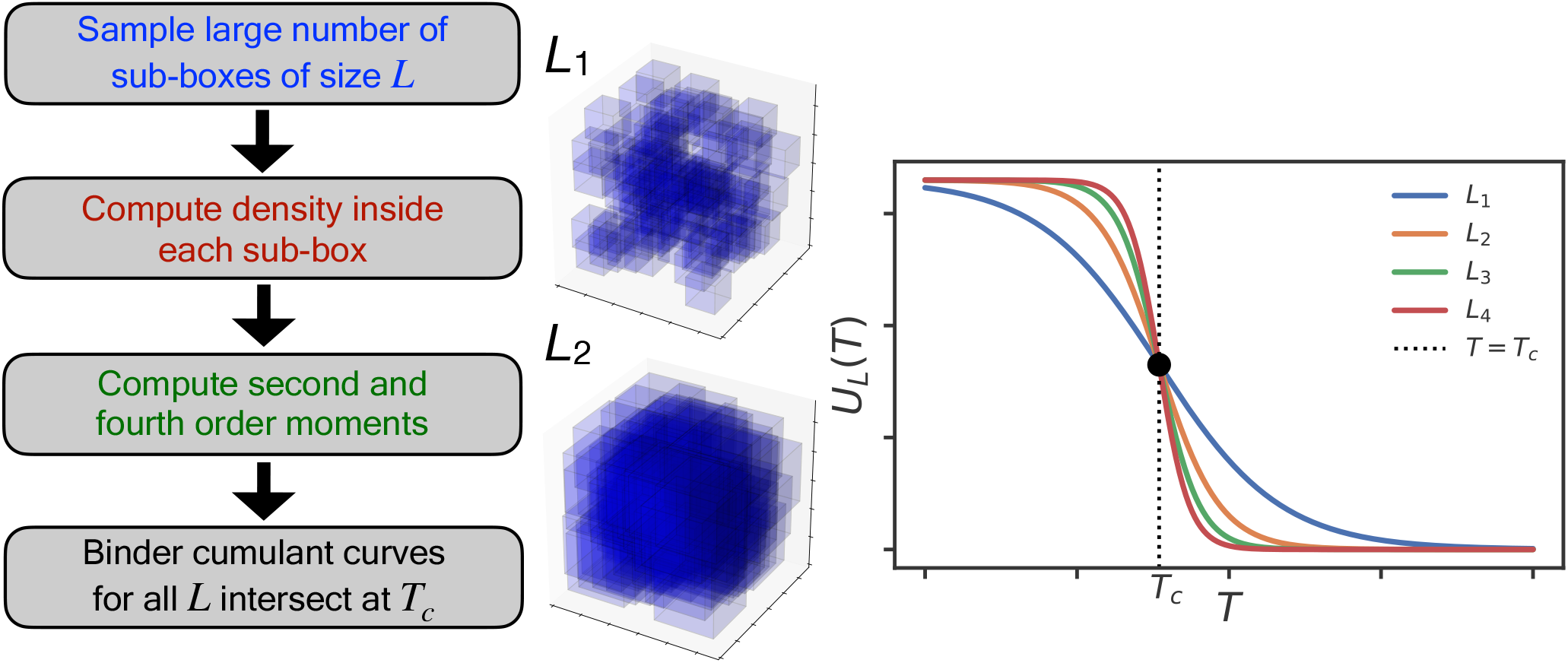
Workflow for Binder Cumulant Analysis. Schematic illustrating the procedure used to obtain an accurate and independent estimate of the critical temperature *T*_*c*_. This is based on computations of higher order moments of the order parameter *m*_*L*_ = (*ϕ*_*L*_(*T* ) − ⟨*ϕ*_*L*_(*T* ) ⟩), which quantifies the deviation of the density from its mean value within a cubic sub-box of length *L* at temperature *T* . A large number of cubic sub-boxes of length *L* are sampled. The variance and the fourth-order cumulant are obtained from *m*_*L*_ for each of the sampled sub-boxes. The curves for Binder cumulants, computed for different *L* values, intersect at *T*_*c*_.

Probability distributions, *p*(*ϕ*), calculated within sub-boxes of different size *L*, are shown in Fig. 9(A-D) for four representative temperatures (also see Fig. S3 in the *SI Appendix*). Below *T*_*c*_, and for small sub-boxes, the density distributions are bimodal Fig. 9(A-C), reflecting the coexistence of dense and dilute phases. The density distributions become unimodal and Gaussian for all sub-box sizes *L* as *T*_*c*_ is approached and crossed (Fig. 9D).

**Fig. 9.**
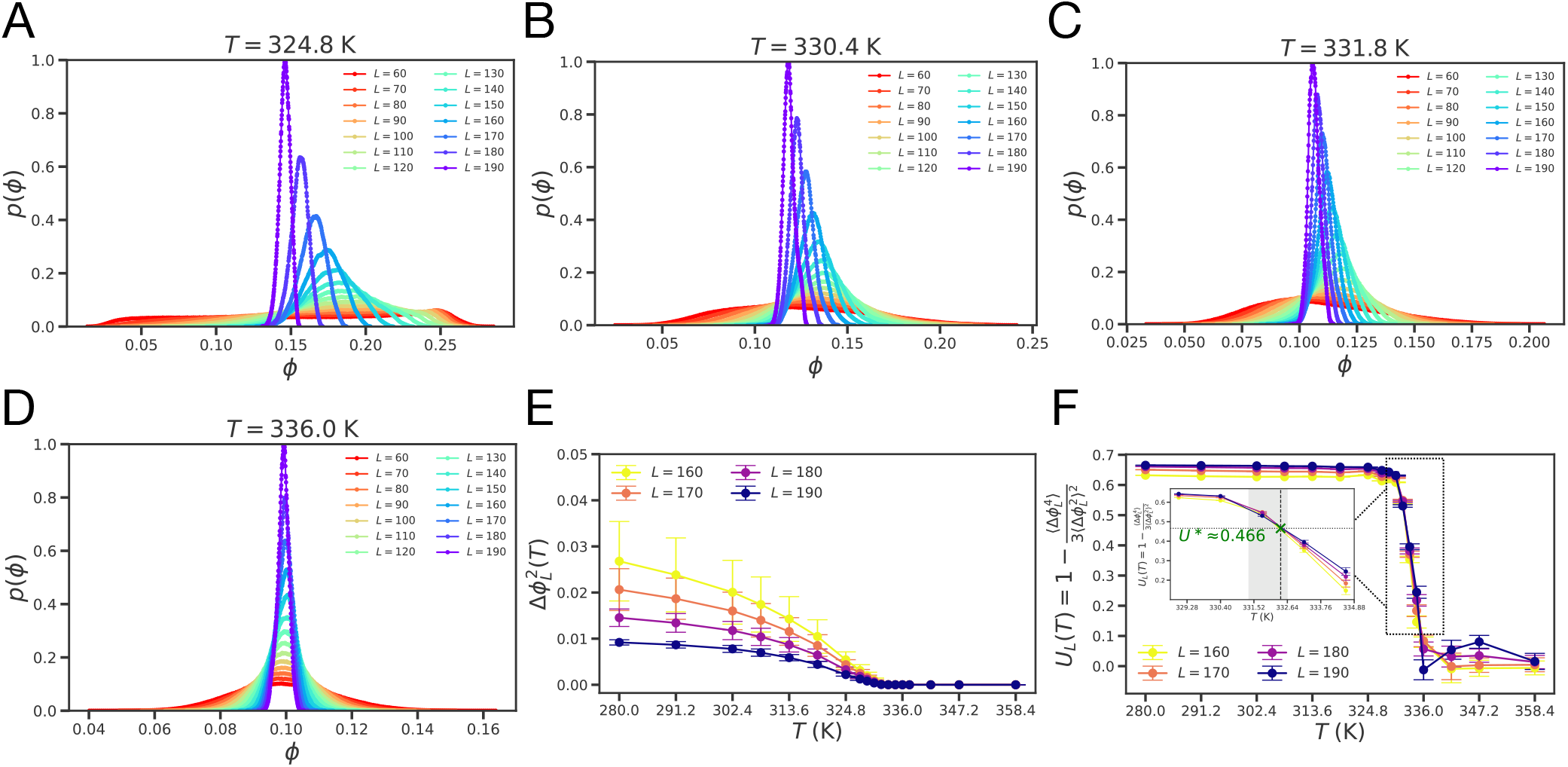
Distributions of the order parameter and results from Binder cumulant analyses. (A–D) Probability distributions of the sub-box densities *p*(*ϕ*) at various temperatures *T* and for different sub-box sizes *L* (in lattice units). These distributions are used to calculate the second- and fourth-order moments of the density fluctuations. Below *T*_*c*_, and for small sub-boxes, the density distributions are bimodal, reflecting the coexistence of dense and dilute phases. The density distributions become unimodal and Gaussian for all sub-box sizes *L* as *T*_*c*_ is approached and crossed. (E) The variance of the order parameter plotted as a function of the temperature *T* for various sub-box sizes *L*. (F) Binder cumulant *U*_*L*_(*T* ) as a function of *T* for various sub-box sizes *L*. The curves intersect at a median critical temperature of *T*_*c*_ ≈ 332.42 K, as shown in the inset. The shaded gray region (in the inset) indicates the range between the lower and upper bounds of the pairwise crossing temperatures, and the green cross marks the intersection point corresponding to the median *T*_*c*_. At the median value of *T*_*c*_, the value we obtain for *U*_*L*_ corresponds to *U*^*^= 0.466, which is the value that has been determined for *U*_*L*_ at the *T*_*c*_ of the 3D Ising model (99).

For each sub-box of length *L*, the variance was computed using 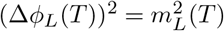 and the fourth-order cumulant was computed using: 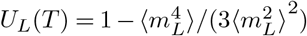, where *m*_*L*_ = (*ϕ*_*L*_(*T* ) − ⟨*ϕ*_*L*_(*T* )⟩) is the order parameter. When the variance and fourth-order cumulant are respectively plotted as a function of *T* for various sub-box sizes *L*, curves of 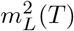 and 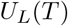 intersect at a common point that corresponds to the true *T*_*c*_ (Fig. 9E, Fig. 9F). Note that this analysis was performed for cumulants computed in sub-boxes of size *L* ≤ 160 l.u. This choice ensures that all density fluctuations are fully accounted for in estimating *T*_*c*_. The mean and median values of *T*_*c*_ were found to be 332.14 K and 332.42 K, respectively (Fig. 9E, Fig. 9F). At the median value, we find that *U*_*L*_(*T* ) ≈ 0.466. This is congruent with the value of *U* ^*^ = 0.466 that is expected at *T* = *T*_*c*_ for the 3D Ising model (99). The statistical error in the estimation of *T*_*c*_ is obtained from the lower and upper bounds of the set of all pairwise crossings of the Binder cumulant curves (shown in the inset of Fig. 9F). The values of these lower and upper bounds are 331.35 K and 332.63 K respectively.

From the point at which the horizontal line that intersects the binodal and the ordinate at *T*_*c*_ one can drop a vertical line and geometrically estimate *ϕ*_*c*_ as the value where this line intersects the abscissa. This approach yields a value of *ϕ*_*c*_ ≈ 0.099 (Fig. 10A). Using this method and the bounds in our estimate of *T*_*c*_, we obtain values of ≈ 0.094 and ≈0.122 as the lower and upper bounds on the estimate of *ϕ*_*c*_, respectively. As an independent approach to estimate *ϕ*_*c*_, we used the law of rectilinear diameters (Fig. 10B). Usage of this law is valid because the interactions are short-range and the system is in 3-dimensions (100, 101). The temperature-dependent average of the dense and dilute phase volume fraction *ϕ*_diam_(*T* ), computed at different temperatures in the near-critical regime, is fit using linear regression and the knowledge of *T*_*c*_. A limitation of the law of rectilinear diameters is its sensitivity to the number of data points used in the regression near *T*_*c*_. To quantify the uncertainty in the estimate obtained for *ϕ*_*c*_, we used bootstrap sampling with replacement of the points in the near-critical regime. For each of the 1000 resampled datasets, we fixed *T*_*c*_ = 332.42 K and fit 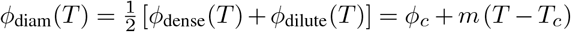, and obtained *ϕ*_*c*_ as the intercept. The bootstrap distribution yields mean and median estimates of *ϕ*_*c*_ ≈ 0.124 and ≈ 0.122, respectively, with a standard deviation of ≈ 0.012 (Fig. 10B). The reported slope, *m* ≈ − 0.003 K^−1^, corresponds to the average value obtained across all the bootstrap samples.

**Fig. 10.**
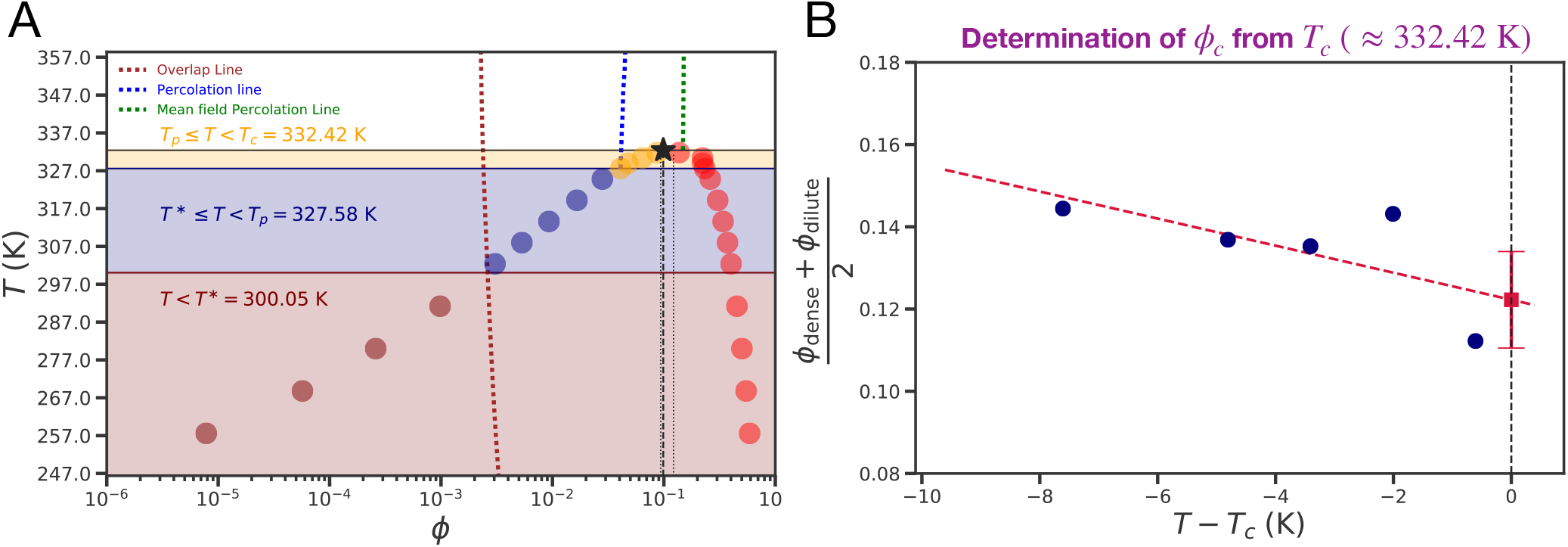
Fully annotated binodal for WT A1-LCD and determination of *ϕ*_c_. (A) Fully annotated binodal for WT A1-LCD. The overlap and the percolation lines are indicated by maroon and blue dashed lines, respectively. The mean field percolation line, mapped using the generalized bond percolation model (21) (see *SI Appendix* ), is shown as a green dashed line. The overlap and the percolation lines intersect the left arm of the binodal and divide it into three distinct regimes. The critical point (*ϕ*_*c*_ ≈ 0.099, *T*_*c*_ ≈ 332.42 K) is marked by an asterisk. Here, *ϕ*_*c*_ is estimated by a vertical linear extrapolation and identified as the value where the vertical drop from *T*_*c*_ intersects the abscissa (shown by the dashed black line). We also show the upper and lower bounds in the estimation of *ϕ*_*c*_ using two dotted black lines that correspond to vertical extrapolations done from the upper and lower bounds in the estimation of *T*_*c*_ (shown in Fig. 9F as the bounds of the shaded gray region). Red circles denote the dense phase points, while the dilute phase is shown by circles whose color corresponds to the regime to which the points belong. These regimes are: (i) the low-temperature regime (maroon), for *T < T*^*^ ; (ii)the intermediate regime (navy blue), for *T*^*^ ≤ *T < T*_*p*_; (iii) the high-temperature regime closest to criticality (orange), for *T*_*p*_ ≤*T < T*_*c*_. (B) An independent estimate of the critical volume fraction *ϕ*_*c*_ was obtained using the law of rectilinear diameters, with *T*_*c*_ = 332.42 K fixed from Binder cumulant analysis. Bootstrap analysis with 1000 resampled datasets yields mean and median estimates of *ϕ*_*c*_ equal to 0.124 and 0.122, respectively, with a standard deviation of 0.012.

Finally, having demarcated the critical and off-critical regimes and determined *T*_*c*_ and *ϕ*_*c*_ separa tely, we analy zed the scaling of the order parameter 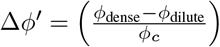 as a function of the distance from the critical point (*ϕ*_*c*_, *T*_*c*_). This is written as: Δ*ϕ*^′^ = *A*_*o*_(1 − *T/T*_*c*_)^*β*^. A regression analysis aimed at simultaneous extraction of *β* and *A*_0_ (58) becomes problematic because of the unavoidable sparseness of the number of points in the critical regime. Additionally, the order parameter is not symmetric around the critical point. Instead, even in the vicinity of the critical point, the change in the volume fraction along the dilute arm is larger than the change along the dense arm, thus making Δ*ϕ*^′^ asymmetrical about *T*_*c*_. Given these considerations, we extracted the value of β by analyzing the ratio 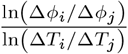 for different pairs of points (*ϕ*_*i*_, *T*_*i*_) and (*ϕ*_*j*_, *T*_*j*_). Here, Δ*ϕ*_*i*_ = (*ϕ*_dense,*i*_ − *ϕ*_dilute,*i*_) and Δ*T*_*i*_ = (*T*_*c*_ − *T*_*i*_). This approach was inspired by the method of Meng et al., that was used to quantify scaling exponents for denatured states of proteins (78). Note that all points (*ϕ*_*i*_, *T*_*i*_) and (*ϕ*_*j*_, *T*_*j*_) were drawn from the critical regime defined by *T > T*_Δ,crossover_. This analysis yielded a set of values for *β*, the mean and median of which were *β*_mean_ = 0.3245 and *β*_median_ = 0.3485, respectively. For the 3D Ising model, *β* = 0.3264 (102). Thus, we conclude that the critical regime of A1-LCD belongs to the same universality class as the 3D Ising model, which is in accord with prior expectations (20, 44, 51, 53–55, 58, 84, 85, 87).

### Locating the percolation line

The percolation threshold, *ϕ*_perc_, is the concentration above which the A1-LCD molecules form a system-spanning network. We computed the threshold concentrations *ϕ*_perc_ for temperatures above and up to *T*_*c*_. The locus of *ϕ*_perc_ values is the percolation line (see Fig. S4 in *SI Appendix*, Fig. 10A). We also used the generalized bond percolation model (21) to compute the mean field percolation line that ignores chain connectivity and correlations (see *SI Appendix*). The mean field percolation line intersects the binodal at *T*_*c*_ (29). However, in accord with previous studies (36, 103), we find that the actual percolation line, computed via rigorous clustering analysis (46, 104), intersects the binodal at a temperature *T*_*p*_ = 327.58 K *< T*_*c*_.

### The overlap line

Next, we quantified the overlap concentration *ϕ*^*^ as a function of *T* . Above the overlap concentration, the likelihood of polymers interacting via intermolecular interactions is higher than the likelihood of them interacting via intramolecular interactions (38, 82). The overlap line is the locus of *ϕ*^*^ values computed as a function of *T* (see Fig. S5 in *SI Appendix*, Fig. 10A). We find that the overlap line intersects the left arm of the binodal at a temperature we designate as *T* ^*^. Thus, for *T* ^*^ ≤ *T < T*_*p*_, the dilute phase is actually a semidilute solution (38, 82, 88) with the chain concentration being above the overlap value. For WT A1-LCD, *T* ^*^ ≈ 300.05 K.

### The binodal has three regimes

The binodal of A1-LCD (Fig. 10A) can be demarcated into three different regimes. These are: Regime I defined by *T < T* ^*^, Regime II defined by *T* ^*^ ≤ *T < T*_*p*_, and Regime III defined by *T*_*p*_ *≤ T < T*_*c*_ (Fig. 10A).

### The dense phase is a percolated network

Across the three regimes of the binodal, the concentration of the dense phase changes by less than a factor of four. Direct calculations of the percolation line and the full binodal show that the right arm of the binodal, which is the locus of *ϕ*_dense_ values, is located to the right of the percolation line. Thus, *ϕ*_dense_ *> ϕ*_perc_ for all *T < T*_*c*_. That *ϕ*_dense_ *> ϕ*_perc_ for all *T < T*_*c*_ clearly indicates that the phase transition of A1-LCD is consistent with PSCP (56, 60).

### Clustering in dilute phases is different across the regimes

In contrast to the dense phase, the concentration of coexisting dilute phase decreases by more than four orders of magnitude between *T*_*c*_ and the lowest temperature for which phase behavior was investigated. Demarcation of the binodal into three regimes derives from the intersection of the dilute arm of the binodal by the overlap and percolation lines. Thus, we reasoned that properties of coexisting dilute phases must differ across the three regimes. To test this hypothesis, we analyzed cluster-size distributions in coexisting dilute phases as a function of the temperature *T*.

The probability *p*(*n*) of forming clusters containing *n* molecules can be described using a power-law relation of the form: *p*(*n*) = *n*^−*τ*^ (105). The magnitude of *τ* is a measure of the extent of clustering. Higher values of *τ* are indicative of smaller clusters being formed, whereas smaller values of *τ* imply the converse. We analyzed cluster-size distributions in the coexisting dilute phases as a function of temperature. Using linear regression analysis, which required the fitting of log10 *p*(*n*) to − *τ* log10 *n*, we extracted the temperature dependence of *τ* (Fig. 11D).

**Fig. 11.**
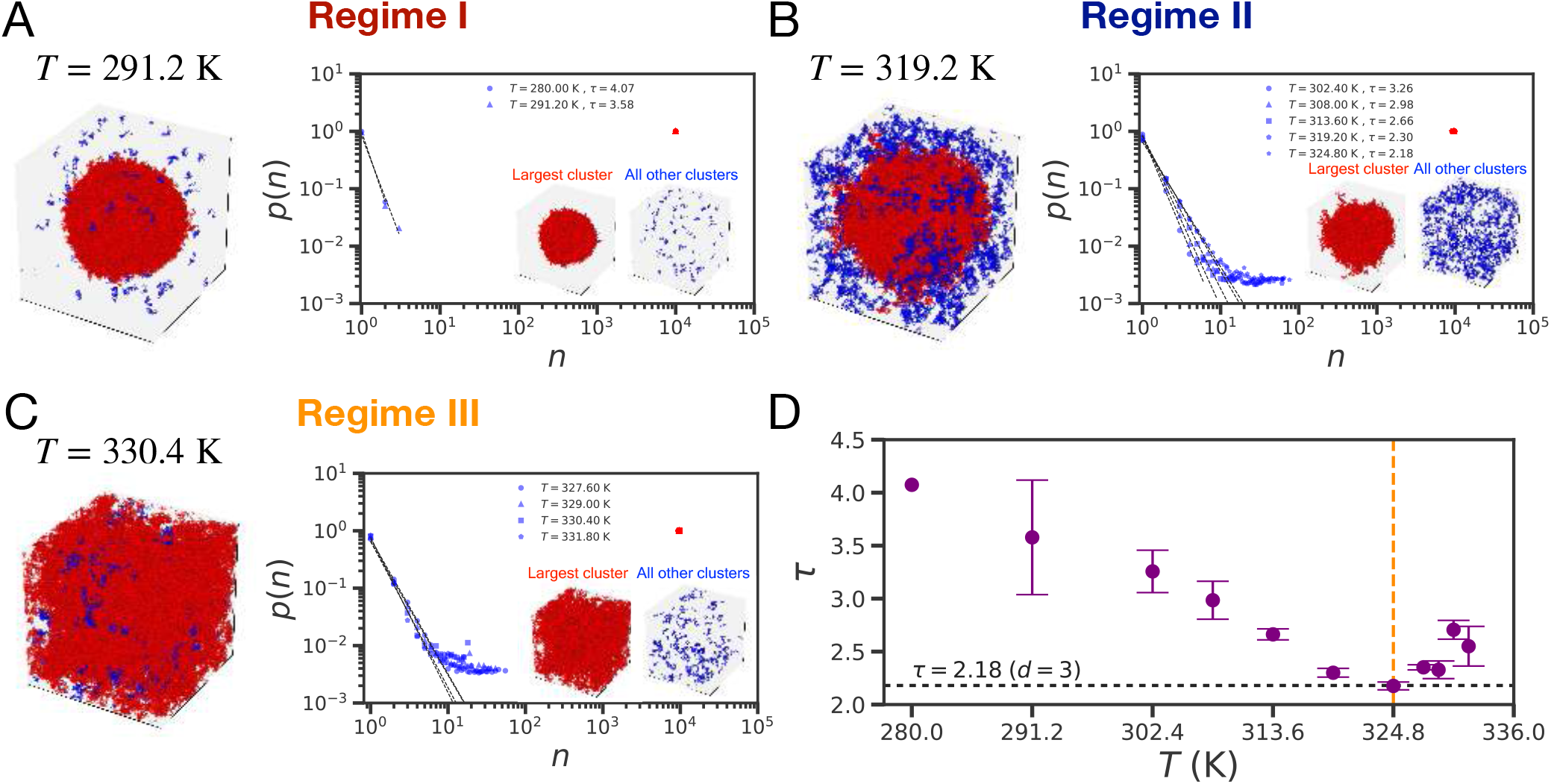
Cluster-size distributions in coexisting dilute phases for each of the three distinct regimes of the binodal. (A-C) Snapshots showing the largest connected cluster (red) and all other clusters (blue), and the cluster-size distributions for the dilute phase (shown by blue symbols) and dense phase (shown by red symbols). The legends show the dilute phase *τ* values obtained at different temperatures *T*, for Regimes I and II. (A) Regime I of the binodal defined by *T < T*^*^. The dense phase that is representative of the largest connected cluster is colored in red, while the dilute phase is shown in blue. (B) Regime II of the binodal defined by *T*^* ≤^ *T < T*_*p*_. Here also, the dense phase that is representative of the largest connected cluster is colored in red, while the dilute phase is shown in blue. (C) Regime III of the binodal defined by *T*_*p ≤*_ *T < T*_*c*_. Here, the snapshot shows the largest connected cluster (red) and smaller clusters (blue) that form. (D) Temperature dependence of the Fisher exponent *τ* for dilute phase cluster-size distributions. The exponent *τ* extracted from the power law fits of the dilute phase cluster-size distributions is plotted as a function of the temperature *T* . The orange vertical line marks *T* = 324.80 K, where *τ* ≈ 2.18, which is the universal critical exponent for three-dimensional percolation (*d* = 3). Above *T*_*p*_, the cluster-size distributions show large-scale fluctuations. As a result, the delineation of dilute and dense phases becomes problematic because they are interconnected. This makes for noisy estimates of *τ* beyond the percolation threshold.

In Regime I, the cluster-size distribution in the coexisting dilute phase decays steeply as *n* increases. The distribution is tailless, and the values of *τ* that we extract lie in the interval 3.6 ≤ *τ ≤* 4.1, with the lower values corresponding to higher temperatures (Fig. 11A). The coexisting dilute phase in Regime I is akin to a “gas” of A1-LCD molecules. In Regime II, the coexisting dilute phase is a semidilute solution (38, 82, 88). Individual A1-LCD molecules can overlap with one another, forming finite-sized clusters in the dilute phase. The cluster-size distributions are characterized by exponents that decrease from *τ* = 3.26 for *T* ≈ *T* ^*^ to *τ* ≈ 2.18 for *T* ≈ *T*_*p*_ (Fig. 11B, Fig. 11D). Across Regime II, the cluster-size distributions have heavy tails with exponential cutoffs.

As the concentration of the coexisting dilute phase increases above the overlap line, cluster formation in the dilute phase becomes easier, as evidenced by the decreasing values of *τ* (Fig. 11D). As a result, larger clusters form, and their abundance, quantified by the values of *p*(*n*), increases with increasing *T* . Additionally, in the cluster size range of ∼50 *< n <* 100, the distributions are characterized by flat tails. These flat tails imply that the probabilities of realizing clusters of different sizes become equivalent for clusters comprising ten to hundred molecules. Similar observations were made by Lan et al., in live cells (49). In addition to the formation of heterogeneous distributions of clusters in the dilute phase, Regime II is characterized by an order of magnitude decrease in the interfacial tension which goes from ≈ 100 *µ*N*/*m at *T* ≈ *T* ^*^ to less than 10 *µ*N*/*m as *T*_*p*_ is approached.

In Regimes I and II, the dense phases can be described as “confined physical gels” (29). This is because *ϕ*_dense_ *> ϕ*_perc_, and the single largest cluster localizes to the dense phase (Fig. 11A, Fig. 11B) across the range of temperatures corresponding to the two regimes. Note that the interfacial tension *γ* is highest in Regime I and decreases to ≈ 10*µ*N*/*m at the highest temperature within Regime II. Thus, capillary fluctuations are expected to increase as the interfacial tension decreases (92). This in turn will weaken the confinement of the percolated network.

Percolation is a continuous networking transition that is different from density transitions (9, 26, 83, 104, 106). It is defined by a distinct critical point that need not coincide with (*ϕ*_*c*_, *T*_*c*_), which is the critical point for density transitions. In 3 dimensions, *τ* = *τ*_*p*_ = 2.18 is the Fisher exponent that defines the percolation threshold (104). *ϕ*_*p*_ = *ϕ*_perc_ is the value of *ϕ* where the percolation line intersects the binodal, and hence (*ϕ*_*p*_, *T*_*p*_) is the critical point for percolation. As *T* approaches *T*_*p*_, we find that *τ* approaches and crosses the Fisher exponent, *τ*_*p*_ = 2.18 (Fig. 11D). The mapping of the percolation line was performed without consideration of the critical exponent that defines the percolation threshold. Thus, analysis of the cluster-size distribution provides independent corroboration of *T*_*p*_ being the temperature where the percolation threshold is crossed.

Taken together, we find that along the dilute arm of the binodal, Regime III (*T*_*p*_ *≤ T < T*_*c*_) is bracketed by two critical points (*ϕ*_*p*_, *T*_*p*_) and (*ϕ*_*c*_, *T*_*c*_). This leads to its designation as the critical regime. In Regime III, both the dense and dilute phases become system-spanning (Fig. 11C). Due to the large density fluctuations and low interfacial tension, the dense phase percolated network is no longer confined. It swells to become system-spanning. This swelling of the dense phase network is concordant with inferences made by Smokers and Spruijt from their analysis of how volumes of dense phases change as the critical point is approached and crossed (45). Above *T*_*p*_, the percolated networks corresponding to the dense and dilute phases swell and become interconnected (Fig. 11C).

### Estimating the theta temperature

The critical point (*ϕ*_*c*_, *T*_*c*_) marks the end of the two-phase regime, implying that the system enters the one-phase regime for *T > T*_*c*_. Thus, for *T < T*_*c*_, the two-body interaction coefficient (*B*_2_) is expected to be negative. *B*_2_ being negative would imply that the effective interactions between pairs of chains or pairs of residues within a chain are attractive for *T < T*_*c*_. The theta temperature (*T*_*θ*_) is the temperature at which *B*_2_ = 0, and therefore for a system such as A1-LCD, which has an upper critical solution temperature (UCST), it follows that *T*_*θ*_ should be greater than *T*_*c*_ (82, 88). Accordingly, we estimated *T*_*θ*_ and asked if it follows canonical expectations.

In Flory-style two-parameter theories (82), *T*_*θ*_ is the temperature where *B*_2_ = 0 for a pair of residues in a chain, and this is taken to be congruent with the temperature at which an individual polymer in a dilute solution behaves like a Gaussian chain (51, 69, 107). To identify the lowest temperature at which Gaussian chain statistics are observed, we performed simulations of individual A1-LCD molecules as a function of the temperature *T* (Fig. 12A). To extract the values of *ν*(*T* ), we analyzed the scaling of ensemble-averaged segmental distances *R*(*s*) plotted against the segment length *s* (Fig. 12B). These plots are referred to as internal scaling profiles (108). For a given profile, *R*(*s*) quantifies the ensemble-averaged spatial separation *R*(*s*) for all linear segments of length *s*. If the scaling of *R*(*s*) against *s* shows power law behavior, then the implication is that *R*(*s*) ∼ *s*^*ν*^. A value of *ν* = 0.5 defines a chain that follows Gaussan statistics.

**Fig. 12.**
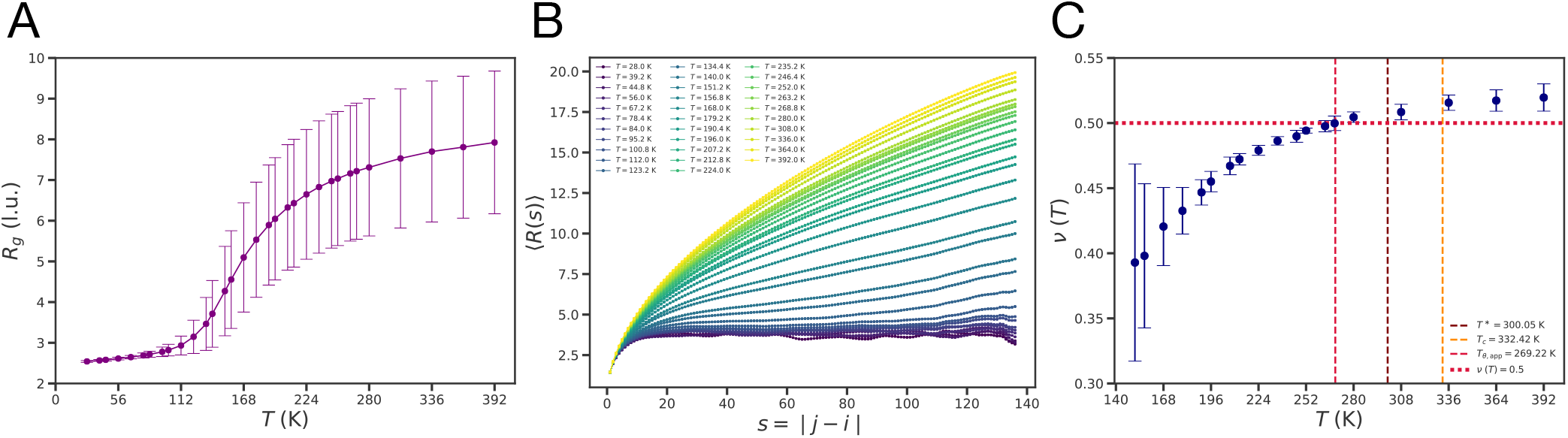
Conventional scaling analysis severely underestimates the theta temperature (*T*_θ_) for A1-LCD. (A) Radius of gyration (*R*_*g*_) values from single chain simulations of A1-LCD as a function of the temperature *T* . These were computed from the last 25% of the snapshots from the equilibrated simulations and across three replicates for each temperature. Error bars denote the standard deviation in the computed *R*_*g*_ for each temperature. (B) *R*(*s*) versus *s* for single A1-LCD chains at various temperatures *T* . (C) *ν* values obtained from analysis of the scaling of *R*(*s*) versus *s* as a function of *T* . The *ν* values were extracted as the slopes of plots of ln(*R*(*s*)) against ln(*s*). This analysis was restricted to temperatures where fractal behavior was evident and not for profiles where *R*(*s*) is essentially flat because this corresponds to the globule regime. The inferred (apparent) theta temperature *T*_*θ*,app_, corresponding to where *ν*(*T* ) = 0.5, is approximately 269.22 K (annotated by the red vertical dashed line). This is well below the critical temperature *T*_*c*_ ≈ 332.42 K. The overlap temperature *T*^*^ and the critical temperature *T*_*c*_ are annotated by maroon and orange vertical dashed lines, respectively. The scaling analysis based on the segmental distances *R*(*s*) as a function of the segment length *s* results in a severely erroneous estimate of the theta temperature for A1-LCD.

From the analysis of the internal scaling profiles extracted at different temperatures, we can identify the lowest temperature at which *ν* = 0.5 (Fig. 12C). In Flory-style mean field theories, this would be taken as the inferred (apparent) theta temperature *T*_*θ*,app_. We find that the lowest temperature (*T*_*θ*,app_) at which *ν* = 0.5 is 269.22 K. Other approaches (109, 110) introduced in the literature yield similarly low estimates of *T*_*θ*_. These estimates of *T*_*θ*_ are incorrect because they are well below the *T*_*c*_ value of 332.42 K. Our findings negate the applicability of the assumptions of Flory-style two parameter theories, which mandate that *ν* = 0.5 at temperatures where *B*_2_ = 0 for a pair of chain molecules.

Given the errors of, and weaknesses inherent to the scaling analysis, we computed *T*_*θ*_ directly by setting it to be the lowest temperature at which *B*_2_ = 0 for a pair of A1-LCD molecules. For this, we computed potentials of mean force (PMFs) *W* (*r*) for a pair of A1-LCD molecules at different temperatures (Fig. 13A). Each PMF quantifies the work done at a given temperature to bring a pair of chains from infinite separation to a distance *r* between their centers-of-mass. The Mayer *f* -function can be written as *f* (*r*) = exp(−*βW* (*r*)) −1 and the integral of −*f* (*r*) yields the excluded volume, which is used to compute *B*_2_ (see Methods section). The normalized *B*_2_ crosses zero at a temperature of 361.73 K. This value is greater than *T*_*c*_ = 332.42 K and appears to be a more reasonable estimate of *T*_*θ*_ (Fig. 13B). That the scaling analysis underestimates *T*_*θ*_ and that the temperature where *B*_2_ = 0 will be higher than the *T*_*θ*,app_ derived from scaling analysis was established by Kholodenko and Freed via conformational space renormalization group calculations (111). Our results are in accord with their calculations, and have implications for drawing inferences regarding solvent quality for disordered proteins, as discussed below.

**Fig. 13.**
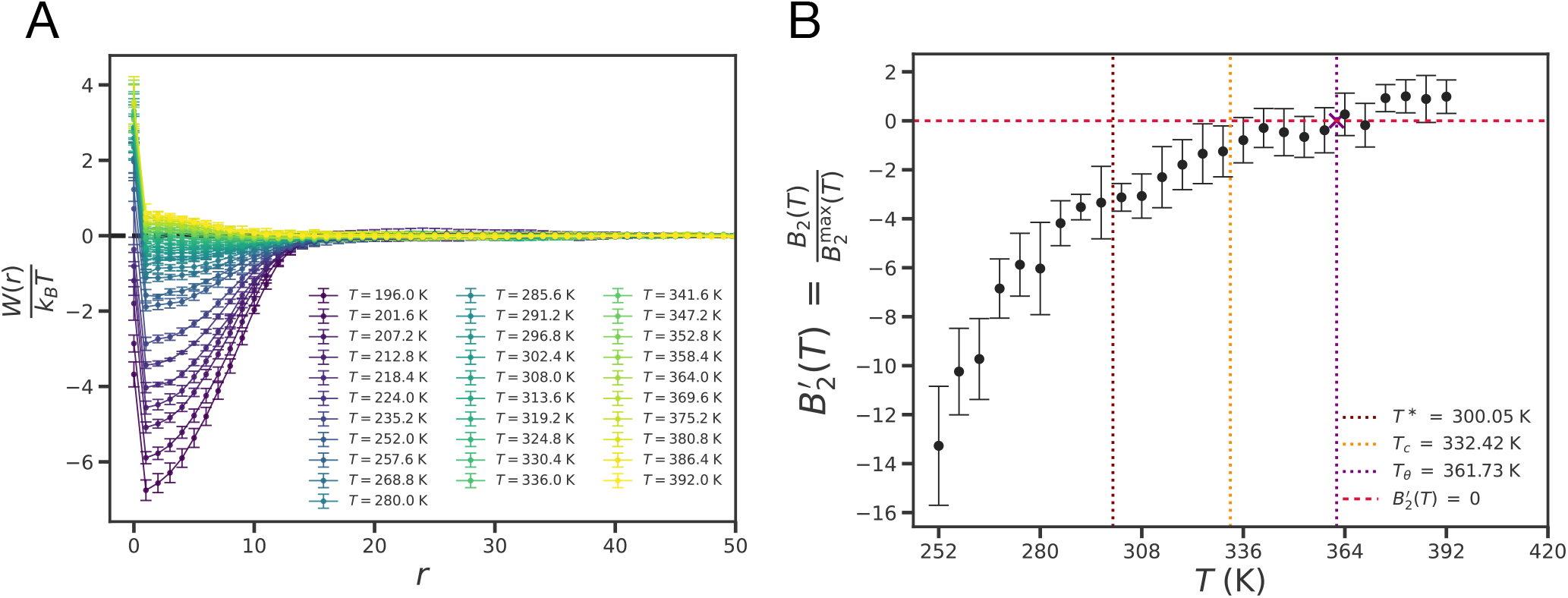
Estimation of the theta temperature (*T*_θ_) for A1-LCD. (A) The potential of mean force *W* (*r*) as a function of the separation between the centers-of-mass of two A1-LCD chains, for various temperatures *T*, extracted from umbrella sampling simulations (see Methods section and Fig. S6 in *SI Appendix* ) using the Weighted Histogram Analysis Method (WHAM) (79–81). For each temperature, umbrella sampling simulations were performed across 6 replicates. (B) The normalized two-body interaction coefficient 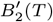 as a function of *T*, obtained from the 2-chain PMF values. The temperature at which 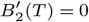 is the theta temperature (*T*_*θ*_). From the temperature dependence of 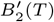, we obtain the theta temperature *T*_*θ*_ using linear interpolation to be ≈ 361.73 K. The overlap temperature *T* theta temperature *T*_*θ*_ are annotated by maroon, orange and purple dotted lines, respectively.

## Discussion

Mapping of critical points is essential for a complete physical description of the phase behaviors of intrinsically disordered proteins such as PLCDs. Knowledge of *T*_*c*_ provides a direct route to obtaining comparative assessments of sequence-specific driving forces for phase transitions (54, 55). In this work, we demonstrate that methods used in the computational literature do a poor job of mapping the critical regime. The main challenge comes from finite size artifacts. Furthermore, incorrect delineation of the mean field versus critical regimes leads to imposition of a single form for the scaling of the order parameter across the entirety of the binodal. Even the analytical form used for the scaling of the order parameter assumes a symmetry that is not present in computed or measured binodals, with there being larger variations in the dilute arm as opposed to the dense arm. We show that these challenges can be overcome by using a systematic and rigorous approach: First, we deploy simulation cells that are large enough, encompassing the appropriate numbers of molecules that allow for collecting statistics regarding densities and fluctuations near the critical point. Second, we delineate the off-critical (mean field) and near-critical regimes by mapping the temperature dependent changes to interfacial widths, from which we derive the interfacial tension. Third, we compute the critical temperature using rigorous finite size scaling approaches based on the analysis of Binder cumulants. This automatically yields an estimate of *ϕ*_*c*_, which we verify independently by applying the law of rectilinear diameters. Finally, with the full binodal in hand, we asked where the overlap and percolation lines intersect the binodal. We find that these lines intersect the left arm of the binodal thereby setting up three distinct regimes for the coexisting dilute phases (36).

Beyond establishing a protocol for accurate mapping of critical points, we uncovered distinct features regarding the cluster-size distributions in coexisting dilute phases that are characteristic to each of the three regimes of the binodal (Fig. 10A, Fig. 11). In a recent study, Varma et al. (112) proposed a connection between the partition coefficient (*K*_*P*_ ) and a normalized temperature distance (*t*) from the critical point. We propose that a more direct assessment of the vicinity to the critical point comes from measurements of cluster-size distributions in coexisting dilute phases. This is feasible in cells (48, 49, 113–115) and in vitro (116–123). When combined with titrations above and below the overlap concentration, cluster-size distributions can be used as diagnostics of proximity to the critical point. For the measurements to be effective, they will need to be able to access the entirety of the cluster-size distribution. While this is challenging for methods that are based on scattering or fluorescence (117, 123), it should be possible using methods such as microfluidic resistive pulse sensing (MRPS) that provide a detection range from 70 nm - 15 *µ*m (116, 121, 122). MRPS has been used to quantify the cluster-size distributions in coexisting dilute phases of different systems (116, 121, 124)

Temperatures are useful proxies for the normalized inter-chain interaction coefficients (*B*_2_) because 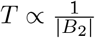 (6, 38, 55). This is relevant for understanding the effects of changes to *B*_2_ on clustering in dilute phases. Lan et al. (49) reported that in unstressed cells, the negative elongation factor (NELF) forms nucleoplasmic clusters that are consistent with what we observe for A1-LCD in the dilute phases of Regime II. Conversely, in stressed cells, the phosphorylation of NELF leads to the formation of macrophases. Setting 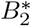 to be the value of the interaction coefficient that corresponds to *T* ^*^, we would propose that 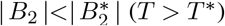 for both unphosphorylated NELF and phosphorylated NELF. At endogenous expression levels, unphosphorylated NELF is in the one-phase regime where it forms clusters corresponding to Regime II. Conversely, phosphorylation moves the system into the two-phase regime by increasing the magnitude of *B*_2_ (making *T* lower), and the coexisting dilute phase is akin to what we observe for Regime II.

Our calculations show that the percolation line intersects the binodal to the left of the critical point. Thus, the dense phase is a confined physical gel defined by an intra-condensate percolated network (9, 24, 25). For A1-LCD and related systems, the networks have been shown to have small-world like topology with smaller-scale density inhomogeneities (56). Quick-freeze deep-etch cryo-electron microscopy of a variant of A1-LCD has provided direct evidence of percolated network-like organization within condensate interiors (116). Additionally, single-fluorogen imaging of A1-LCD condensates (125) has revealed the inhomogeneous organization of molecules within A1-LCD condensate interiors (125).These properties give condensates their tunable viscoelasticity, leading to their designation as viscoelastic network fluids (76, 126–130).

In Regime III, the dense phase becomes unconfined and this network swelling has been inferred via measurements of volumes and volume fractions by Smokers and Spruijt (45). Signatures of being in Regime III will be the formation of interconnected networks of coexisting dense and dilute phases. This behavior has been observed for the WNK1 kinase that forms condensates in response to osmotic stress (131). The observed behavior was referred to as spinodal decomposition. Indeed, Regime III is expected to coincide with the spinodal. Thus, the dynamics of phase separation in Regime III will likely follow that of viscoelastic phase separation (132). System-spanning networks that are consistent with behaviors in Regime III have also been observed in signaling systems that form at membranes (133).

Availability of the full binodal allowed us to assess the accuracy of using internal scaling profiles to identify the theta temperature. We find that this yields an erroneously low estimate for *T*_*θ*_ and the false result of *T*_*c*_ being higher than *T*_*θ*_. Direct computation of *B*_2_ provides a more reliable estimate of *T*_*θ*_ that is in accord with canonical expectations (82). The central tenet of Flory-style two parameter theories is that *ν* = 0.5 for a single chain when *B*_2_ = 0 for pairs of residues (beads) (82). It is known that this is not formally valid for *d* = 3, where *d* is the dimensionality of the system (134). The Flory model comes about by assuming an equivalence between an individual chemical or Kuhn monomer - residue (bead) in our case - and the entire chain. However, the effective two-body interaction coefficient for pairs of chains will be influenced by correlations that stem from chain connectivity, chain flexibility, and three-body as well as higher-order interactions (110). The impact of these correlations leads to a decoupling of the impact of solvent quality on chain size versus shape, which will be pronounced for finite-sized heteropolymers *viz*., IDPs (109, 135).

Our results have broad implications for studies of IDPs. It is customary to use either simulations or measurements to derive internal scaling profiles and quantify the apparent value of *ν* (17, 69, 108, 136–140). The derived *ν* value is then used to judge the quality of the solvent for the IDP of interest. Our results suggest that these inferences of solvent quality must be treated with caution. This is because it is quite likely that the temperature or equivalent control parameter at which *ν* = 0.5 will lie below *T*_*c*_ or the critical value of control parameter. Therefore, any rigorous assessment of solvent quality will require direct computation or measurement of the two-body interaction coefficient *B*_2_, or alternatively, computation or measurement of phase boundaries (60, 141) as a function of temperature or changes to solvent via chemical perturbations such as changes in pH, denaturant, or salt concentration.

Finally, we note that the findings reported here should also be accessible via off-lattice MD simulations. For each temperature, the simulations will need to be on the order of milliseconds, and the simulation cells will need to encompass *O*(10^4^) molecules. A recent study (77) showed that structures within simulated dense phases of A1-LCD molecules share similarities across three different simulation engines that include LaSSI, MPIPI-recharged (142), and CALVADOS (90, 143). Based on these comparisons, we propose that the findings reported are likely to be consistent across different simulation protocols that use similar, single-bead-per-residue models. The key to mapping critical points is the avoidance of finite size artifacts and the use of finite size scaling methods (59, 89). It might also be essential to revisit the use of slab geometries that are in vogue across many MD simulation protocols (20, 51, 53–55, 84, 90, 144). Slabs cause a smoothening of density profiles and this is achieved by suppressing fluctuations that are the defining hallmarks of critical regions. These geometries are helpful deep inside the mean field regime, but are likely to be problematic in the critical regime. Assuming finite size artifacts are avoided and cubic boxes that do not break symmetries *a priori* are used, then the extraction of cluster-size distributions across different temperature regimes should be feasible using MD simulations that can be deployed using different engines.

## Acknowledgements

We are grateful to Tanja Mittag for collaborations, and to Michael Rubinstein for pedagogy and insights.

## Author contributions

Conceptualization: GM, SG, GC, RVP; Methodology: GM, GC, SG, RZ, RVP; Formal Analysis: GM, GC, SG, KMR, RVP; Investigation: GM, SG, RVP; Software: GM, GC, RZ, KMR; Project Administration: RVP; Data Curation: GM, SG, KMR; Visualization: GM, SG, KMR, RVP; Writing - Original Draft: RVP; Writing - Editing and Revising: GM, SG, KMR, RVP; Funding Acquisition: RVP.

## Supporting Information (Appendix) for

### Distinguishing near-versus off-critical phase behaviors of intrinsically disordered proteins

#### Derivation of the percolation threshold from mean-field predictions ϕ_perc,MF_(*T*)

To compute the percolation threshold from a mean field model, we used the generalized bond percolation model of Choi et al., (21). In this model, the mean field connectivity parameter between stickers of types ‘*i*’ and ‘*j*’ is defined as:

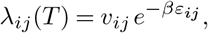

where *v*_*ij*_ is the effective bond volume (configurational volume for forming an *i*–*j* contact), *ε*_*ij*_ is the pairwise interaction energy in units of *k*_B_*T*, and *β* = 1*/k*_*B*_*T* is the inverse temperature. The total connectivity *D* arising from all sticker types is given by (21),

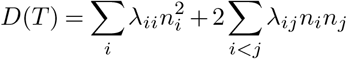

where *n*_*i*_ is the number of stickers of type ‘*i*’ per chain. The percolation threshold is calculated using

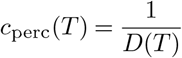

For WT A1-LCD, the relevant sticker types are Y (tyrosine), F (phenylalanine), and R (arginine) (17, 29, 60, 75), where R interacts primarily with the aromatic residues. Expanding *D*(*T* ) gives

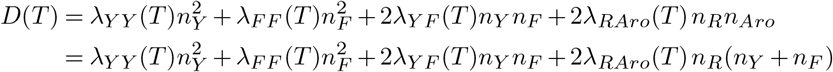

Hence,

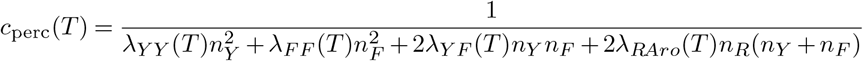

If each chain consists of *N* beads and each bead occupies a volume *v*_*b*_ (*v*_*b*_ = 1 l.u.^3^ in LaSSI units), then the percolation volume fraction is

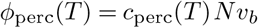

Substituting *c*_perc_(*T* ) gives the final expression:

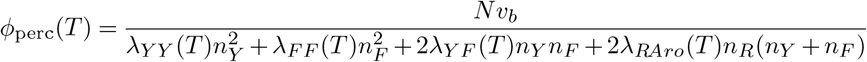

Now,

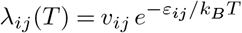

Since *k*_*B*_ = 1 in LaSSI (46, 56), the percolation threshold explicitly depends on temperature as

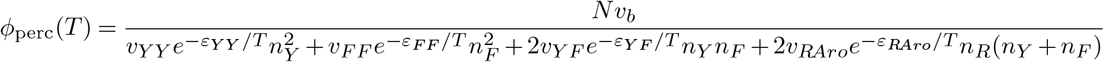

If all bond volumes are taken as being equal (*v*_*ij*_ = 1 l.u.^3^) and *ε*_*ij*_ are the LaSSI contact energies, the final form of the expression is

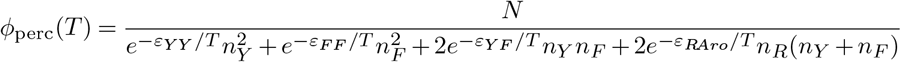

For the WT A1-LCD sequence (N = 137 residues), the relevant sticker types and their counts per chain are:

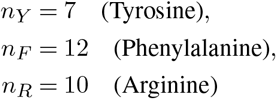

**Fig. 1.**
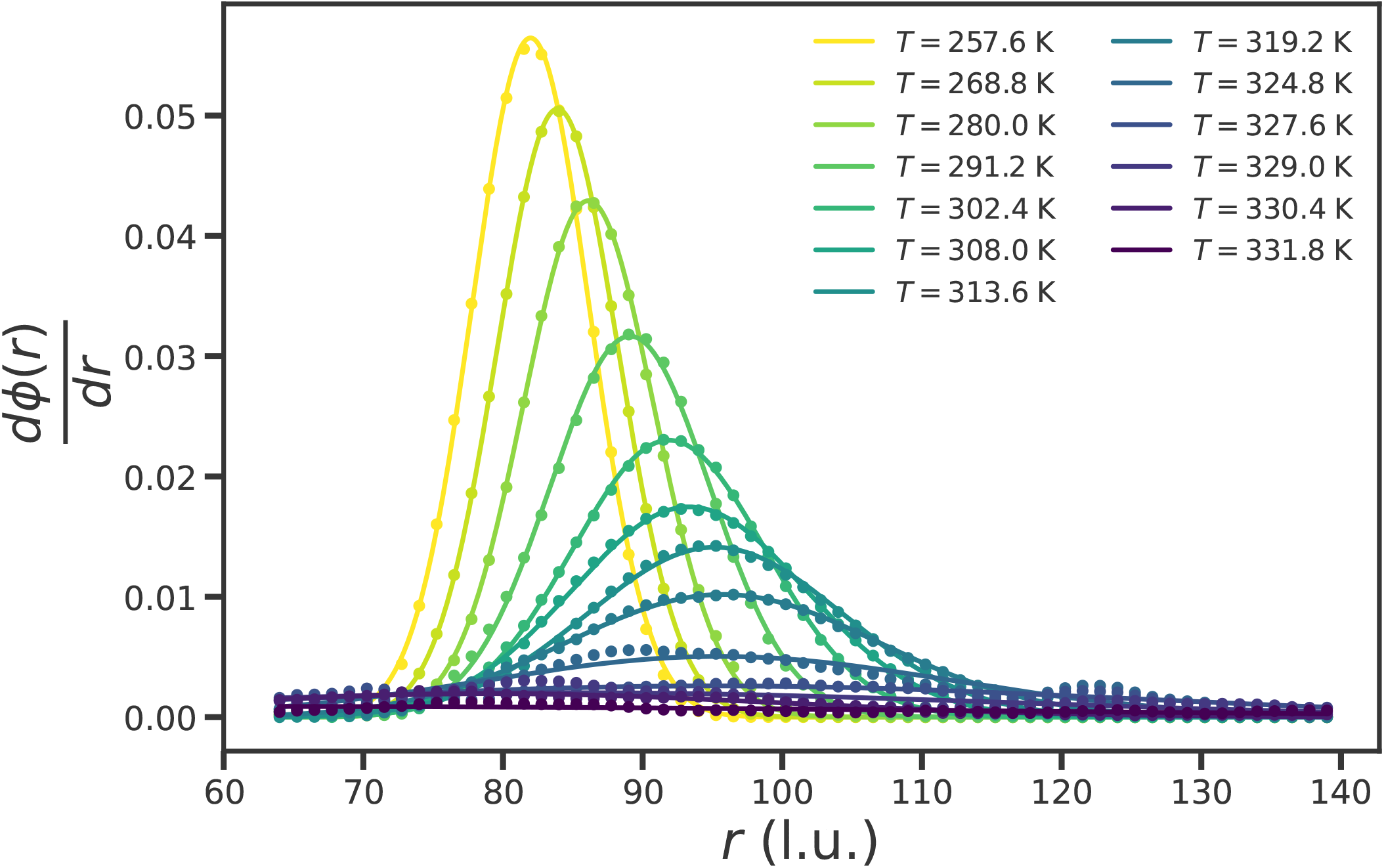
Derivatives of the radial density profiles, *dϕ*(*r*)*/dr*, at various temperatures for the 10^4^-chain system of WT A1-LCD. The derivative of the radial density profile at each given temperature is fit to a Gaussian function (72), from which the interfacial width Δ(*T* ) is extracted as the full-width-at-half-maximum (FWHM), given by 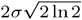, where *σ* is the standard deviation. The midpoint of the interface *r*_mid_(*T* ) is the mode of the Gaussian.

**Fig. 2.**
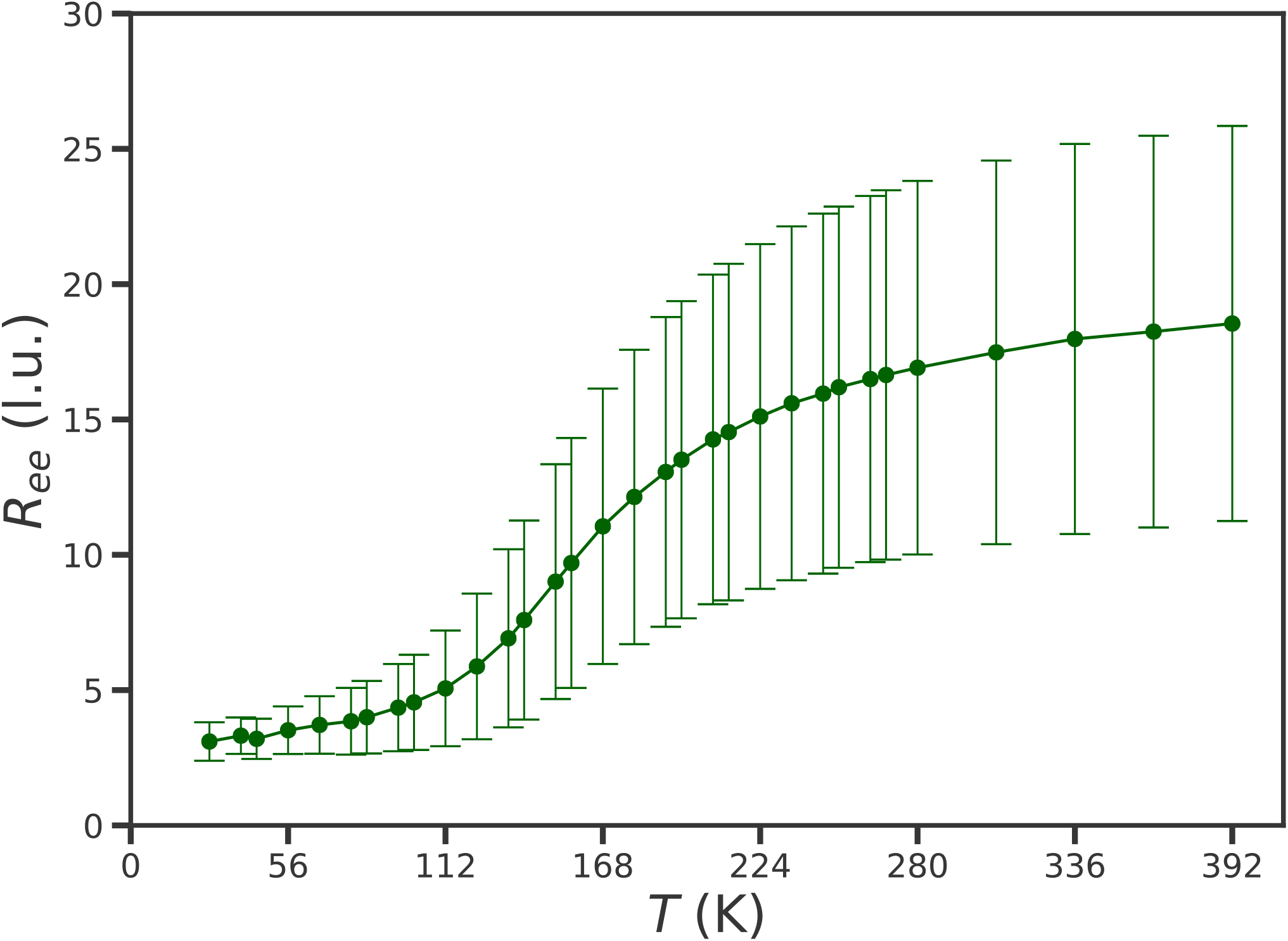
End-to-end distance *R*_ee_ of a single A1-LCD chain (*N* = 137) as a function of the temperature *T*. These were computed from the last 25% of the snapshots from the equilibrated single chain simulations and across three replicates for each temperature. The mean *R*_*ee*_ values across the set of temperatures studied range from ≈ 3 l.u. to ≈ 17 l.u.. Error bars denote the standard deviation in the computed *R*_*ee*_ for each temperature.

**Fig. 3.**
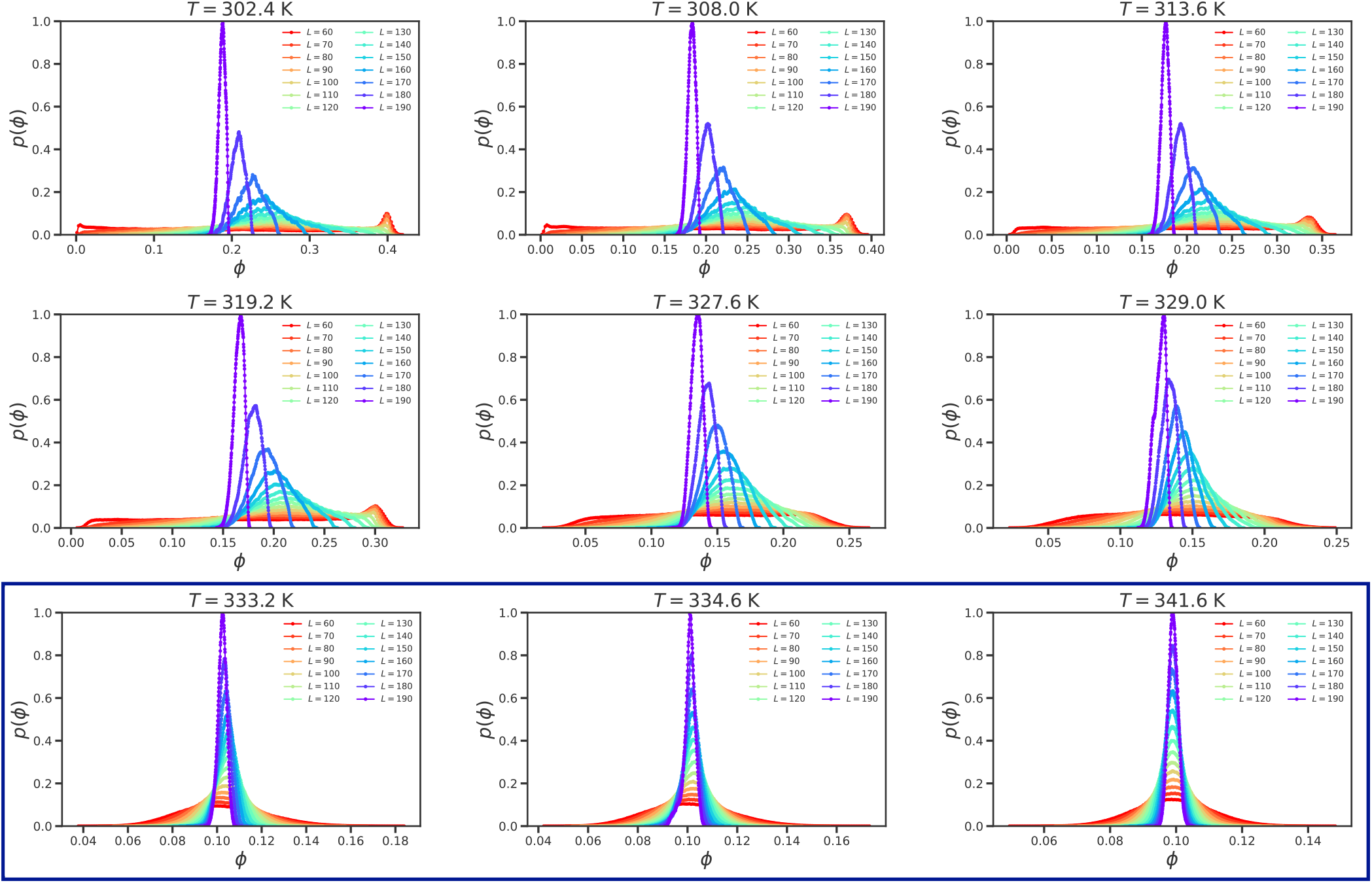
Distributions *p*(*ϕ*) as a function of *ϕ* obtained by sampling sub-boxes of various sizes, at different temperatures. Note that as *T* approaches *T*_*c*_ ≈ 332.42 K, the density distributions show Gaussian behavior for large *L*. In contrast, at lower values of *T* (for which the system is in the two-phase regime) and small sub-box sizes, we observe a bimodality in the density distributions. For each of the temperatures shown inside the navy blue box that corresponds to *T > T*_*c*_, all the distributions collapse onto a unimodal Gaussian form across all sub-box sizes, consistent with homogeneous single-phase behavior.

**Fig. 4.**
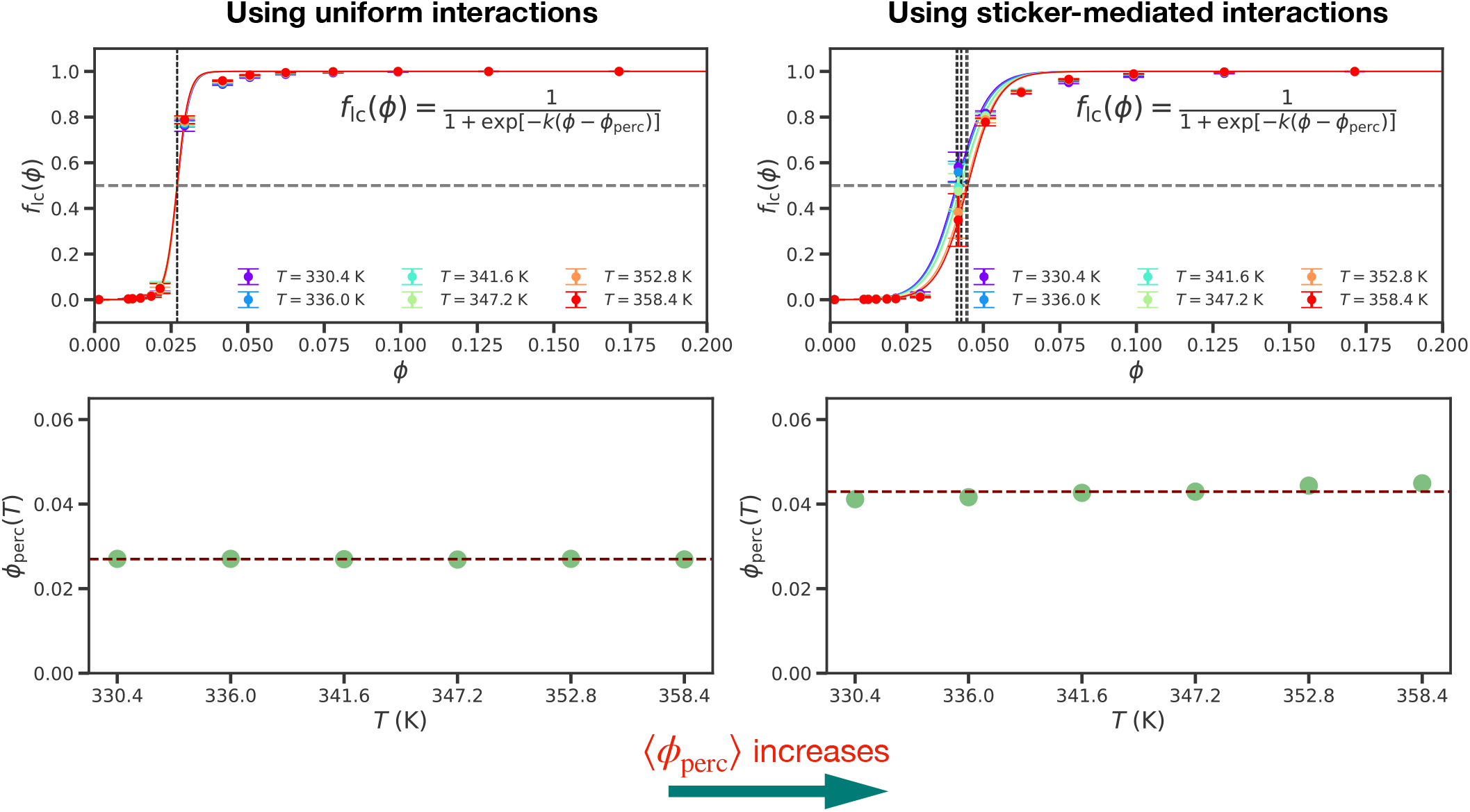
Calculation of the percolation threshold (*ϕ*_perc_(*T* )) using different ways of defining clusters. This calculation uses information regarding the fraction of molecules in the largest cluster 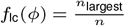, which we compute as a function of the simulation volume fraction *ϕ*. Here, *n* = 10^4^ (the total number of A1-LCD molecules in the system). This analysis is performed for temperatures above and up to the critical temperature *T*_*c*_ for the cases when uniform interactions between all residues are considered, and when sticker-sticker interactions between only the strongest stickers tyrosine (Y), phenylalanine (F) and arginine (R) residues are taken into account. Percolation threshold (*ϕ*_perc_(*T* )) as a function of the temperature *T* (in Kelvin) is also shown for the two cases, and the average *ϕ*_perc_(*T* ) increases for the latter.

**Fig. 5.**
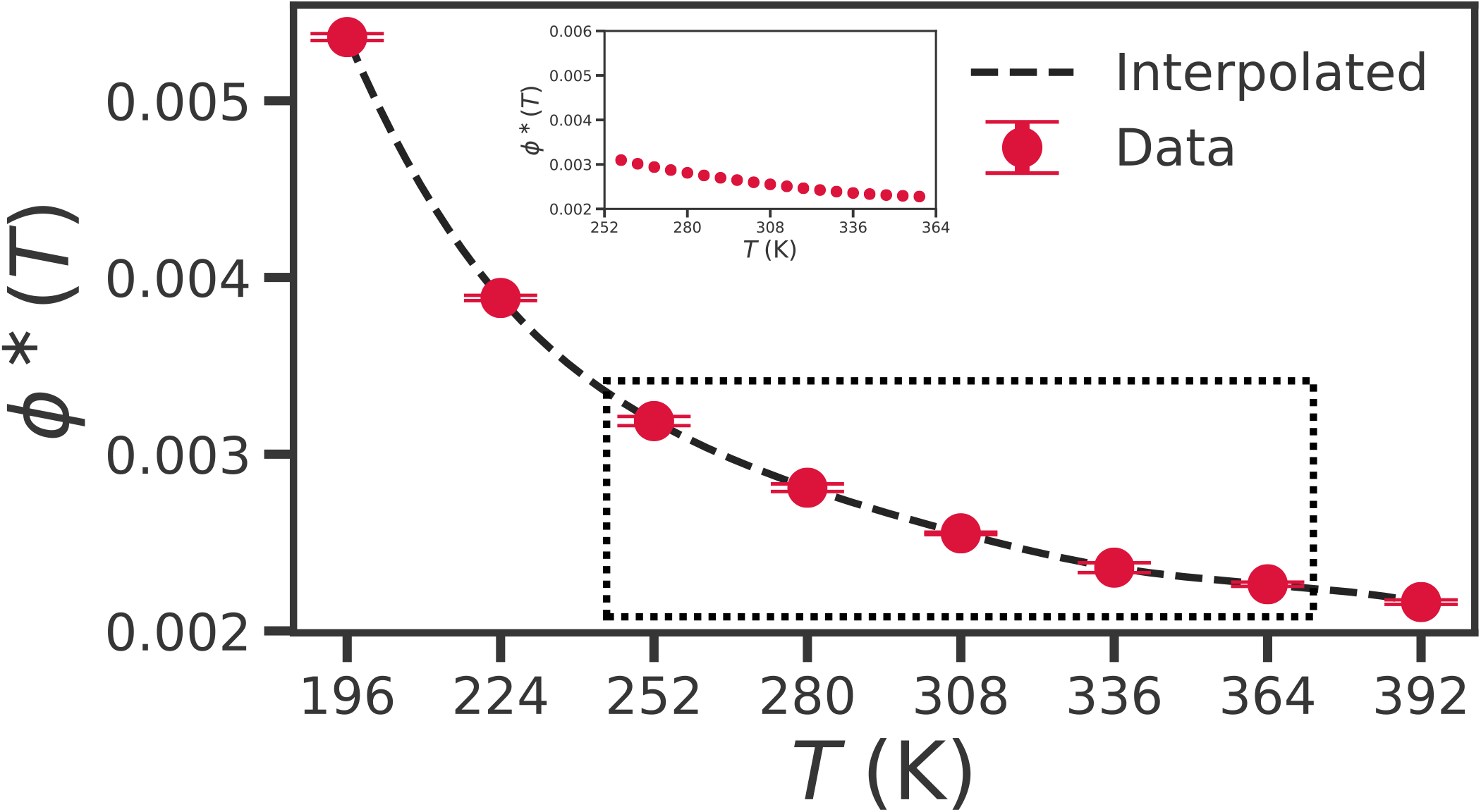
Overlap volume fraction *ϕ*^*^ as a function of the temperature *T*. The data are shown here for the wild-type A1-LCD. The overlap volume fraction was calculated from single chain simulations. The inset shows the data over a zoomed-in range of *T*.

**Fig. 6.**
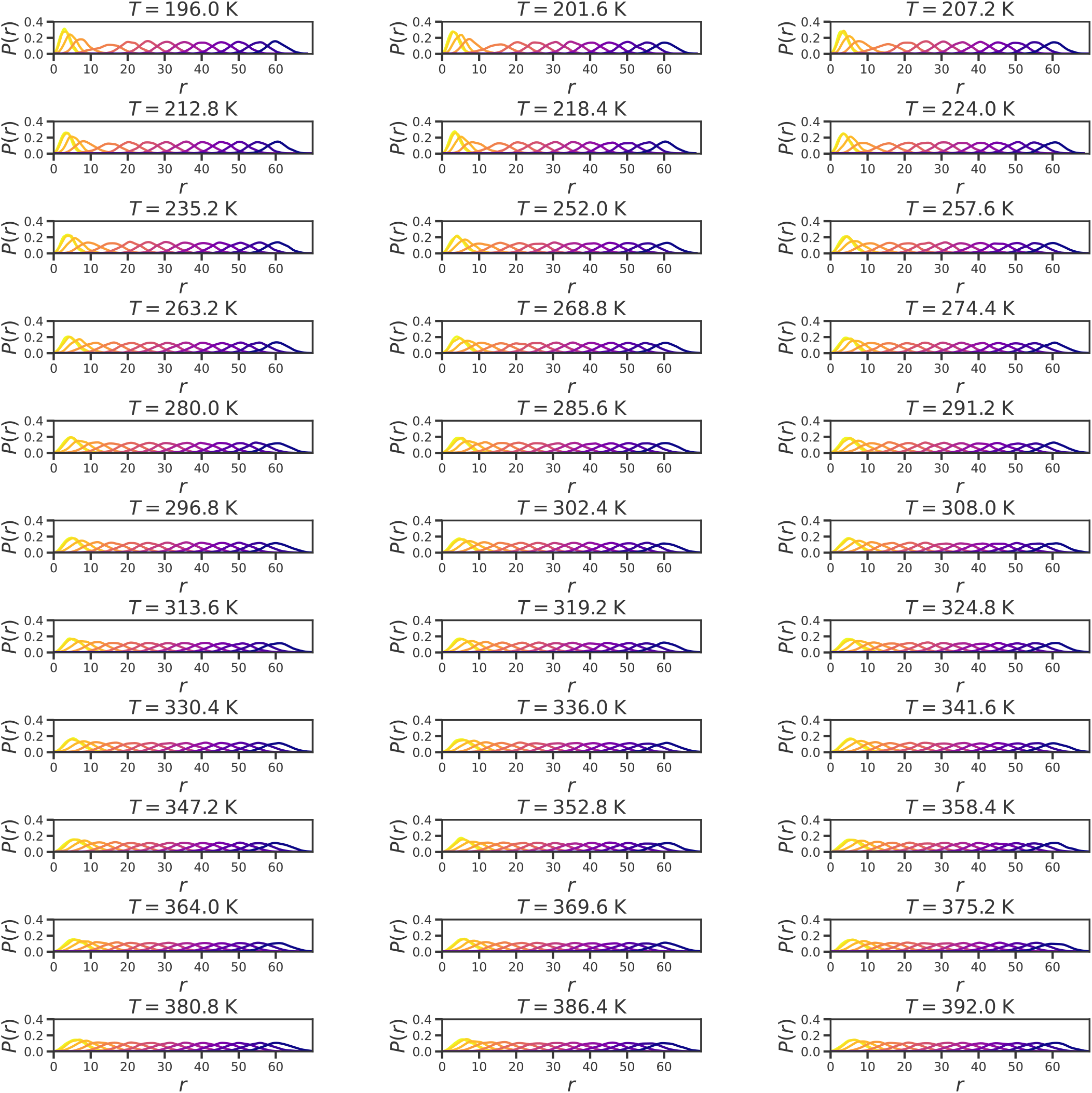
Probability distributions for A1-LCD obtained from two-chain umbrella sampling simulations at different temperatures. Probability distributions *P* (*r*) of the center–of–mass separation *r* between two A1-LCD chains obtained from umbrella sampling simulations performed at different temperatures. Each panel corresponds to a distinct temperature, and colored curves denote individual umbrella windows centered at increasing restraint distances. The spring constant *k*_spring_ was chosen to ensure sufficient overlap between neighboring windows thereby ensuring reliable reconstruction of the unbiased potential of mean force (PMF) using WHAM (79–81). At low temperatures, the distributions are concentrated at small separations, consistent with strong intermolecular association, whereas increasing temperature shifts probability weight toward larger separations, reflecting progressive weakening of attractive interactions.

